# Isotonic and minimally invasive optical clearing media for live cell imaging *ex vivo* and *in vivo*

**DOI:** 10.1101/2024.09.13.612584

**Authors:** Shigenori Inagaki, Nao Nakagawa-Tamagawa, Nathan Huynh, Yuki Kambe, Rei Yagasaki, Satoshi Manita, Satoshi Fujimoto, Takahiro Noda, Misato Mori, Aki Teranishi, Hikari Takeshima, Yuki Naitou, Tatsushi Yokoyama, Masayuki Sakamoto, Katsuhiko Hayashi, Kazuo Kitamura, Yoshiaki Tagawa, Satoru Okuda, Tatsuo K. Sato, Takeshi Imai

## Abstract

Tissue clearing has been widely used for fluorescence imaging of fixed tissues, but not for live tissues due to its toxicity. Here we develop minimally invasive optical clearing media for fluorescence imaging of live mammalian tissues. Light scattering is minimized by adding spherical polymers with low osmolarity to the extracellular medium. A clearing medium containing bovine serum albumin (SeeDB-Live) is minimally invasive to live cells, allowing for structural and functional imaging of live tissues, such as spheroids, organoids, acute brain slices, and the mouse brain *in vivo*. SeeDB-Live minimally affects the electrophysiological properties and sensory responses of neurons. We demonstrate its utility for widefield imaging of subcellular voltage dynamics, such as backpropagating action potentials, in acute brain slices. We also utilize SeeDB-Live for widefield voltage imaging of dozens of dendrites *in vivo*, demonstrating population dynamics. Thus, SeeDB-Live expands the scale and modalities of fluorescence imaging of live mammalian tissues.

---

Live biological tissues are dynamic by nature. They continuously change their morphology as well as the activity of various extracellular and intracellular signaling. Thanks to various chemical and genetically encoded fluorescent biosensors, we can image and measure dynamic biological phenomena within the live tissues and organs using fluorescence microscopy. However, the imaging depth is often limited by tissue opacity. Therefore, imaging the dynamics of the entire biological system is a challenge.

The opacity of the biological tissues is largely due to the inhomogeneity of refractive index within the samples. Two-photon microscopy uses a near-infrared excitation laser instead of visible light to reduce light scattering; however, the imaging depth is limited to a few hundred microns in mammalian tissues *in vivo* ^1^. Adaptive optics uses a deformable mirror or spatial light modulator to correct aberrations caused by macroscopic refractive index distortions, but is not effective for highly scattering samples ^2^. For fixed tissues, optical clearing is a powerful approach: light refraction and scattering are minimized by removing high index components (e.g., lipids) and/or by immersing the sample in high index solutions with refractive indices of 1.43-1.55 ^3–13^. However, most of the clearing agents developed for fixed tissues are toxic to live cells. Less toxic chemicals (e.g., glycerol, dimethyl sulfoxide, sugars, tartrazine, etc.) have been tested for highly fibrous extracellular structures, such as skin and skull *in vivo* ^14–19^; however, these chemicals interfere with cellular functions and/or are hypertonic, hampering their applications to live tissues for imaging of cellular functions. Osmolarity of the clearing agent is particularly critical for live imaging.

Some of the chemicals have been proposed for live cell imaging. Iodinated contrast agents were attractive candidates because of their low osmolarity. One of them, indixanol, has been shown to improve the transparency of bacteria and some multicellular organisms ^20,21^. However, its toxicity to mammalian cells has not been fully evaluated. Another study attempted to improve transparency of the mouse brain by adding glycerol to drinking water ^22^. However, it was not demonstrated whether the refractive index in the brain was changed by feeding glycerol; glycerol should be easily metabolized once absorbed in the gut. Therefore, the optical clearing strategy has not been effectively used for fluorescence imaging of cellular functions (e.g., recording intracellular Ca^2+^ and other signals) in live mammalian tissues.

Here we developed SeeDB-Live, a tissue clearing medium for live mammalian cells and tissues. SeeDB-Live contains bovine serum albumin (BSA), which is minimally invasive to cells and has exceptionally low osmolarity when dissolved in water. SeeDB-Live improved the imaging depth of spheroids, organoids, and brain slices *ex vivo* up to ∼2-fold with confocal and two-photon microscopy. SeeDB-Live is particularly useful for widefield voltage imaging of mammalian brain *ex vivo* and *in vivo* with confocal and widefield imaging, expanding the scale and modalities of fluorescence imaging in the nervous system.

## Results

### Strategies for minimally invasive optical clearing of live mammalian cells

It is well known that light scattering in tissues is caused by refractive index mismatch between the light scatterer and the medium. Previously, simple immersion-based clearing agents (refractive index, 1,47-1.52) have been developed for fixed tissues (e.g., SeeDB with fructose and SeeDB2 with iohexol)^8, 9^; however, osmolarity of these clearing agents are extremely high. To make live tissues transparent in a non-invasive way, we would have to use either i) membrane-permeable or ii) membrane-impermeable low osmolarity (i.e., high molecular weight) chemicals to reduce the refractive index mismatch (Fig. 1a). For i) membrane-permeable chemicals, we do not need to change the concentration of the saline; however, when ii) membrane-impermeable chemicals are added to the medium, we need to subtract the concentration of the saline to keep the medium isotonic. We have listed membrane-permeable (e.g., dimethyl sulfoxide (DMSO), glycerol, propylene glycol) and membrane-impermeable high molecular weight chemicals (e.g., iodinated contrast agents, straight polymers, and spherical polymers) as candidates. Candidate chemicals also need to be highly soluble in water. These chemicals demonstrated a concentration-dependent increase in refractive index when dissolved in water (Fig. 1b).

**Figure 1.**
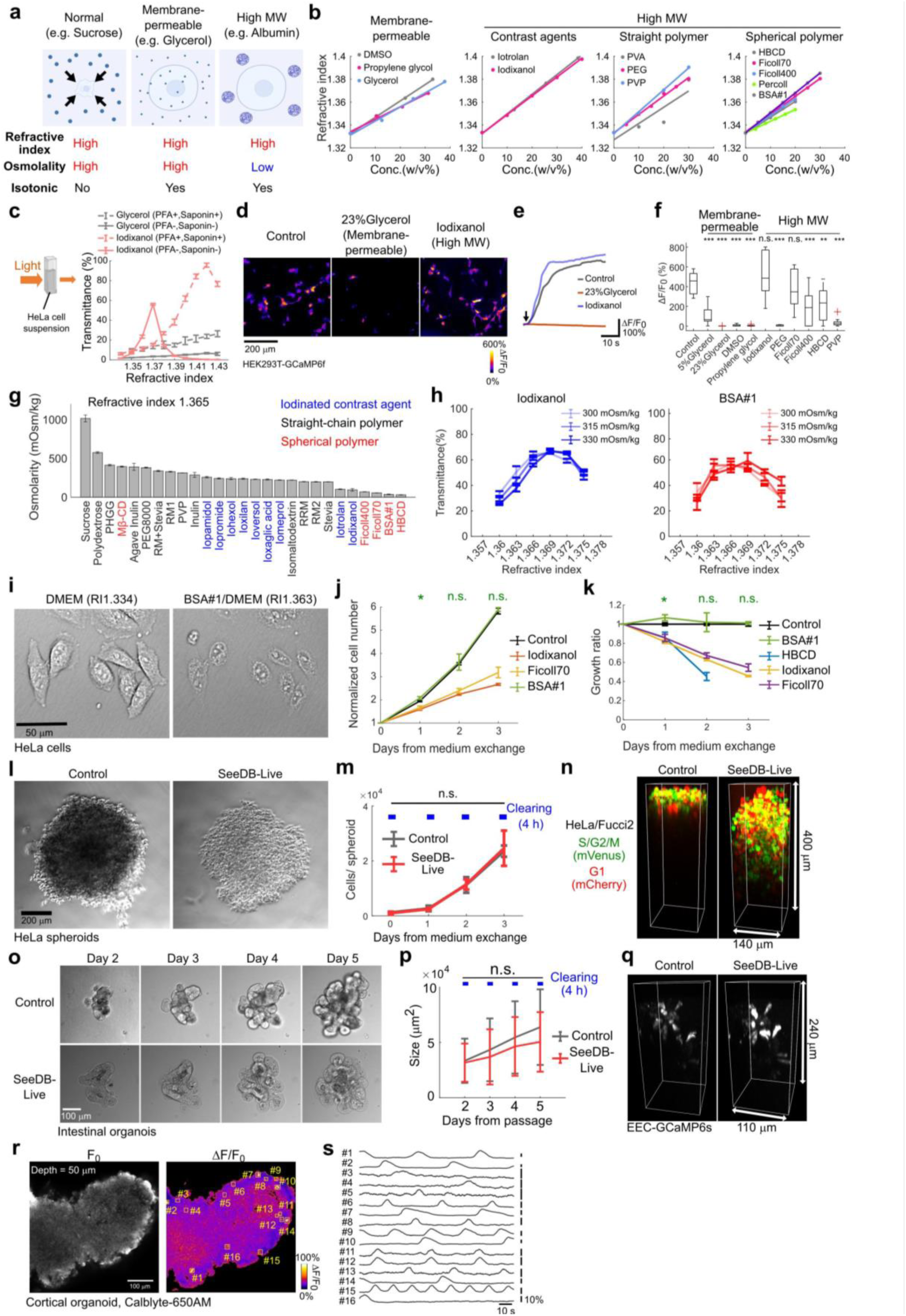
Screening for non-toxic optical clearing agents for imaging live mammalian cells and tissues. **(a)** Principles for optical clearing of live cells with membrane-permeable and membrane-impermeable (high molecular weight) chemicals. **(b)** Concentration-dependent increase in refractive indices with candidate chemicals. **(c)** Transmittance of HeLa cell suspension (4 × 10^6^ cells/mL, transmittance at 600 nm) in isotonic saline solution with glycerol (membrane-permeable) or iodixanol (membrane-impermeable and high molecular weight) at different refractive indices. Fixed cells were treated with PFA and saponin. **(d-f)** Calcium response of GCaMP6f-expressing HEK293T cells to 50 μM ATP stimulation. These cells were tested after 2 hours of culture in a medium containing various candidate chemicals. The refractive index of the medium was adjusted to 1.365 (except for 5% glycerol) by adding these chemicals. Osmolarity was not adjusted to isotonicity. Calcium responses (ΔF/F_0_) **(d)** and time series data **(e)** of cells incubated in normal medium, glycerol medium, and iodixanol medium. ATP was added at the timing indicated by the arrow. Calcium responses in various media are summarized in **(f)**. n = 9 cells from 3 dishes. \*\*\**p* < 0.001; \*\**p* < 0.01; n.s., non-significant (*p* ≥ 0.05) (Dunnett’s multiple comparison test). See also Supplementary Fig. 1a using a plate reader-based assay. **(g)** The osmolality of candidate high molecular weight chemicals in aqueous solution (refractive index 1.365, in ddH_2_O, n = 3 each). Sucrose was used as a control. For many high molecular weight chemicals, the measured values were higher than the theoretical values (e.g., PVP). In contrast, the osmolality of BSA, HBCD, and Ficoll70 was close to the theoretical value. The BSA#1 represent one product of BSA. For other products, see Supplementary Fig. 1c. **(h)** The optimal refractive index of the extracellular medium was determined in PBS at different osmolalities. Transmittance of unfixed HeLa cell suspensions (4 × 10^6^ cells/mL) was measured. Left: optimal refractive index (refractive index 1.369) of iodixanol-containing PBS. Right: optimal refractive index (refractive index 1.363-1.369) of BSA-containing PBS (n = 3 each, see Supplementary Fig. 1d for other wavelengths). **(i)** Phase contrast images of HeLa cells. Left, normal medium; right, BSA-containing medium (refractive index 1.363). **(j)** Proliferation curve of HeLa/Fucci cells in iodixanol, Ficoll70, and BSA#1 containing medium (refractive index 1.363, 320 mOsm/kg). Cell numbers were measured by fluorescence imaging of cell nuclei (n = 5 wells). \**p* < 0.05; n.s.; non-significant (*p* ≥ 0.05) (Dunnett’s multiple comparison test). P-values are <0.001 unless otherwise mentioned. **(k)** Growth ratio in refractive index-optimized (refractive index 1.363, 320 mOsm/kg) and normal medium. Cell counts were measured by fluorescence images of cell nuclei divided by the control at each stage (n = 5 wells). \**p* < 0.05; n.s., non-significant (*p* ≥ 0.05) (Dunnett’s multiple comparison test). P-values are <0.001 unless otherwise mentioned. For other BSA product, see Supplementary Fig. 1e. **(l)** Phase contrast images of HeLa/Fucci2 cell spheroids. Normal medium (left) and SeeDB-Live medium (refractive index 1.366, 320 mOsm/kg; right). Spheroids were cultured in a normal medium and the medium was replaced with SeeDB-Live medium 1 hour before the observation. **(m)** Growth curve of HeLa/Fucci2 cell spheroids with and without treatment with SeeDB-Live for 4 hours per day. Dissociated cells were counted with a hemocytometer (n = 3 spheroids each). n.s., not significant (*p* ≥ 0.05) (Wilcoxon rank sum test corrected with Holm-Bonferroni correction). **(n)** Three-dimensional fluorescence images of a HeLa/Fucci2 cell spheroid in the control and SeeDB-Live media. See also Supplementary Video 1. **(o)** Phase contrast images of intestinal organoids treated with SeeDB-Live (refractive index 1.363) for 4 hours per day. Organoids were embedded in Matrigel. **(p)** The growth of the intestinal organoid culture. The size of organoids (areas in the phase contrast images) in control and SeeDB-Live are shown. Organoids were detected using Cellpose. n = 34, 45, 57, and 63 organoids for control; n= 33, 55, 58, and 56 organoids for SeeDB-Live. n.s., not significant (*p* ≥ 0.05) (Wilcoxon rank sum test corrected with Holm-Bonferroni correction). **(q)** 3D fluorescence images of GCaMP6s-expressing enteroendocrine cells before and after SeeDB-Live treatment in the intestinal spheroid sample. GCaMP6s is specifically expressed in ePet-Cre; Ai162 mice (EEC-GCaMP6s). Normal medium (left) and SeeDB-Live (refractive index 1.366; right). **(r)** Basal fluorescence (left) and ΔF/F_0_ images (right) of a cortical organoid labeled with a calcium indicator, Calblyte-650AM. The images were taken by confocal microscopy. **(s)** Spontaneous calcium transients of neurons indicated in **(j).** MW, molecular weight; DMSO, dimethyl sulfoxide; PVA, polyvinyl alcohol; PEG, polyethylene glycol; PVP, polyvinyl pyrrolidone; HBCD, hyperbranched cyclic dextrin.

Next, we sought to determine the optimal refractive index for clearing live mammalian cells. For this purpose, we prepared a suspension of live or paraformaldehyde (PFA)-fixed and membrane-permeabilized HeLa cells (4 × 10^6^ cells/mL) in isotonic solution of iodixanol in phosphate buffered saline (PBS); to keep the osmolarity of the buffer isotonic, we mixed isotonic iodixanol solution (60% w/v) and PBS to prepare isotonic solutions with different refractive indices. It is known that refractive indices used to clear fixed tissues are typically 1.43-1.55 ^10^. Indeed, the PFA-fixed and membrane-permeabilized HeLa cells were most transparent at a refractive index of ∼1.42. In contrast, we found that the live HeLa cells are most transparent at a refractive index of ∼1.37, much lower than the fixed cells (Fig. 1c). Moreover, the optimal range of the refractive index was relatively narrow for live cells. These results indicate that the refractive index of live cells (cytosol) is ∼1.37, and index matching between cytosol and extracellular medium (typically 1.33-1.34) will dramatically improve the transparency of the tissues. We also tested a membrane-permeable chemical, glycerol, at a wide range of refractive indices. While glycerol was known to clear skin and skull^17^, it was not effective at all for clearing live cells (Fig. 1c).

We next examined whether intracellular functions remain intact in the presence of candidate chemicals. Using GCaMP6f calcium sensor, we evaluated the calcium responses of HEK293T cells to 50 μM ATP solution under various clearing media at refractive index 1.365 (e.g., 23.0% w/v in buffer for glycerol) (Fig. 1d-f, Supplementary Fig. 1a, b). Calcium responses were completely abolished in the presence of membrane-permeable chemicals, glycerol, dimethyl sulfoxide, and propylene glycol, while a lower concentration of glycerol (5%) showed weaker responses. These results indicate that membrane-permeable clearing agents greatly impairs cellular functions even at refractive index 1.365. Among the membrane-impermeable, high molecular weight chemicals, straight polymers abolished calcium responses (e.g., polyvinyl alcohol and polyvinyl pyrrolidone). In contrast, intact calcium responses were observed for iodinated contrast agents (e.g., iodixanol) and spherical polymers (e.g., Ficoll70). These results indicate that some of the membrane-impermeable, high molecular weight chemicals would be useful for index matching of the extracellular medium without compromising cellular functions.

### Screening and optimization of low osmolarity and non-toxic clearing agents

Membrane-impermeable chemicals will not directly interfere with intracellular functions but may increase osmolarity. To keep the clearing medium isotonic, we have to reduce the concentration of the saline; however, the extracellular ionic conditions would indirectly affect cellular functions. Therefore, the ideal chemical should have a low osmolarity when dissolved in water to achieve the optimal refractive index. The increase in osmolarity can be minimized if we use high molecular weight chemicals (>1 kDa); however, extremely large particles (>10 nm scale) will cause Rayleigh scattering. Straight-chain polymers had prohibitively high osmolalities, much higher than the theoretical values based on molar concentrations ^23^; in contrast, we found that spherical polymers of higher molecular weight, such as bovine serum albumin (BSA), highly branched cyclic dextrin (HBCD), and Ficoll70, have low osmolalities at a refractive index of 1.365 (Fig. 1g, Supplementary Fig. 1b).

The osmolarity of the human serum is typically 285 ± 5 mOsm/L. The osmolarity of the saline buffers and culture media for mammalian cells ranges from 230 to 340 mOsm/L, but is typically within 300-330 mOsm/L. For both iodixanol and BSA, we further refined the optimal refractive index for this range. We prepared PBS at 300, 315, and 330 mOsm/kg with refractive indices of 1.360-1.375 using iodixanol or BSA. The highest transparency was found at refractive index 1.363-1.369 when prepared at 300-330 mOsm/kg (Fig. 1h, Supplementary Fig. 1d). At refractive index 1.366 and 315 mOsm/kg, the concentration of PBS had to be reduced to 71.2% and 80-90 % (e.g., 81.2% for BSA, product #1), respectively, indicating that more physiological ionic condition is maintained when BSA is used. The best refractive index was higher at higher extracellular osmolarity, likely because cytosol is more condensed. When live HeLa cells were incubated with BSA-containing medium (refractive index 1.363), the plasma membrane was almost invisible under the phase contrast microscopy (Fig. 1i).

Using index-optimized isotonic culture medium, we evaluated long-term toxicity using HeLa cells. Twenty-four hours after seeding HeLa cells in plastic dishes, the medium (DMEM) was replaced with the index optimized isotonic DMEM. Cell growth was monitored for up to three days. Cell growth was comparable to the control DMEM for one of the BSA products (BSA #1) (Fig. 1j, k, Supplementary Fig. 1e, f). However, cell growth was lower for iodixanol, highly branched cyclic dextrin, Ficoll70, and some of the BSA products (Fig. 1j, k, Supplementary Fig. 1e, g).

BSA is also preferable in terms of lower viscosity (Supplementary Fig. 1h, i) and specific gravity (Supplementary Fig. 1j). Lower viscosity results in more efficient diffusion of the clearing media. Culture with iodixanol is difficult because the specific gravity of the cell is lower than that of the iodixanol solution and cells easily detach and float in the medium ^20^. Other proteins may be similarly useful; however, BSA is highly soluble in water and is one of the most affordable proteins. In addition, albumin is the most abundant protein in the serum and has been widely used for the mammalian cell culture media (as a major component of fetal bovine serum), suggesting that BSA is minimally invasive to the mammalian cells.

It is known that albumin buffers divalent cations (e.g., Ca^2+^ and Mg^2+^) ^24^. We, therefore, optimized the concentration of Ca^2+^ and Mg^2+^ in the media to keep the concentrations of free divalent cations physiological (Supplementary Fig. 2); the optimal concentration was ∼2-fold higher than the conventional buffer based on the evaluation in neurons (i.e., total concentrations of Ca^2+^ and Mg^2+^ were 3 mM higher). Primary cultures of mouse cardiomyocytes (Supplementary Fig. 3a-c) and hippocampal neurons (Supplementary Fig. 3d-g) were maintained in the BSA-containing culture medium for at least three days without any obvious signs of toxicity.

In this way, we established BSA-containing clearing media for live mammalian cells, named SeeDB-Live, with optimal refractive index (1.363-1.366), osmolality (230-330 mOsm/kg), and Ca^2+^/Mg^2+^ concentrations (2-4 mM each) with saline or culture medium (80% concentrations of normal saline/media) (Supplementary Table 1).

### Fluorescence imaging of live spheroids and organoids with SeeDB-Live

We examined whether SeeDB-Live is useful for fluorescence imaging of multicellular structures. We cleared cultured HeLa/Fucci2 cell spheroids ^25^ with SeeDB-Live (Fig. 1l). Growth of HeLa/Fucci2 spheroids was slightly slower when continuously cultured in SeeDB-Live, possibly due to lower circulation of oxygen (Supplementary Fig. 4a) ^26^. However, daily clearing with SeeDB-Live for 4 hours per day did not affect the growth of the spheroid culture (Fig. 1m). In the confocal microscopy, mVenus and mCherry signals were visible up to ∼100 μm in depth in the control medium; in contrast, signals were visible up to ∼250 μm in the SeeDB-Live medium with confocal microscopy (Fig. 1n, Supplementary Fig. 4b-d, Supplementary Video 1). The brightness of the signals was improved in the deeper area of the spheroids, and the number of detected cells was increased when cleared with SeeDB-Live (Supplementary Fig. 4d).

Next, we tested SeeDB-Live for imaging intestinal organoid cultures. Intestinal organoids are monolayered epithelia, and a luminal (apical) membrane is formed inside the organoid when cultured in the Matrigel. Intestinal organoids were developed from ePet-Cre; Ai162 mice, in which enteroendocrine cells express a calcium indicator, GCaMP6s. The intestinal organoids became transparent after the incubation in SeeDB-Live (Fig. 1o), and the organoid growth was not affected by daily 4-hour clearing with SeeDB-Live (Fig. 1p). The luminal cavity of the organoid was less transparent (Fig. 1o), suggesting that BSA does not efficiently penetrate the tight junctions formed by the epithelial tissues. Nonetheless, the GCaMP6s-positive enteroendocrine cells were visible in deeper areas under SeeDB-Live using confocal microscopy (Fig. 1q, Supplementary Fig. 4e). Calcium imaging demonstrated robust responses to high potassium stimulation, indicating that their physiological functions are maintained (Supplementary Fig. 4f). We also tested SeeDB-Live for confocal calcium imaging of the neuroepithelial and cortical organoids induced from mouse ES cells (Fig. 1r, s, Supplementary Fig. 4g-j) ^27^. Thus, SeeDB-Live will be useful for functional assays of organoids.

### Clearing acute brain slices with SeeDB-Live

Volume imaging is in high demand for neuroscience applications. We evaluated the performance of SeeDB-Live using acute brain slices from Thy1-YFP-H mice (Fig. 2a). After the recovery of acute brain slices in oxygenated artificial cerebrospinal fluid (ACSF), confocal and two-photon images were acquired (Fig. 2b, c). The brain slices containing the cerebral cortex were then perfused with SeeDB-Live/ACSF for 1 hour and imaged again under the same conditions. The imaging depth was increased ∼2-fold for both confocal and two-photon microscopy under SeeDB-Live (Fig. 2c-e, Supplementary Video 2, 3). Similar results were obtained for the hippocampus (Supplementary Fig. 5a-c, Supplementary Video 4, 5). The optimal refractive index of SeeDB-Live was ∼1.366 in acute brain slices (Supplementary Fig. 5d), consistent with our results for cultured cell data (Fig. 1h).

**Figure 2.**
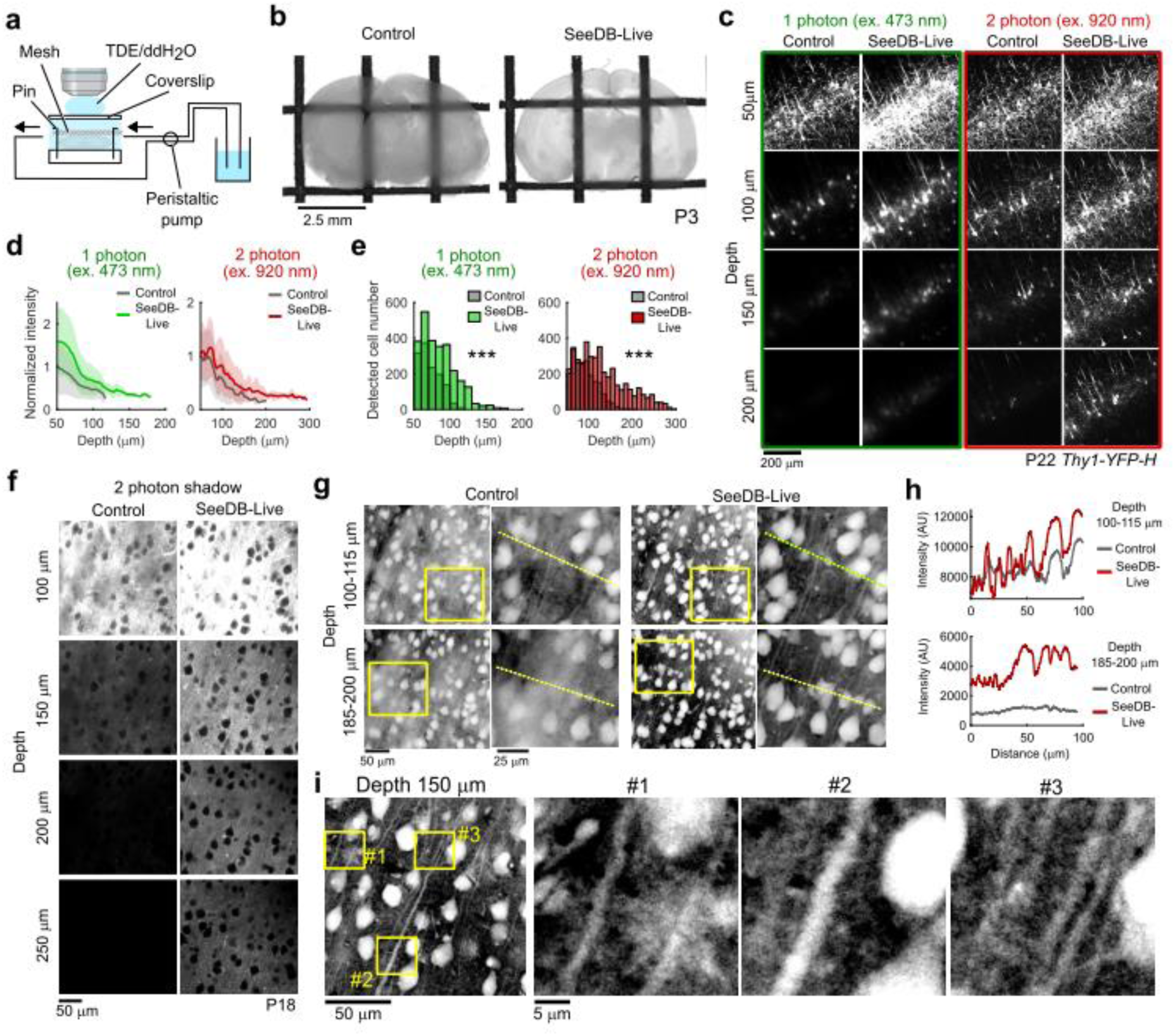
Improved morphological imaging of acute brain slices with SeeDB-Live. **(a)** Preparation of acute brain slices and clearing with SeeDB-Live. Perfusion chamber (1.5 mL/min) was used to immerse acute brain slices with SeeDB-Live. **(b)** Acute brain slices (300 μm thick, age P3, transmission images) before and after clearing with SeeDB-Live. **(c)** *X-y* fluorescence images of an acute brain slice of S1 L5 pyramidal cells. Confocal (1P) and two-photon (2P) images are shown. P22 Thy1-YFP-H mice were used. See also Supplementary Video 2 and 3. **(d)** Fluorescence intensity from cell bodies in *x-y* fluorescence images of S1 L5 pyramidal neurons. ROIs were generated by Cellpose, and mean fluorescence in ROIs are shown. **(e)** Number of detected cells in each depth in S1. Cells were detected by Cellpose. \*\*\**p* < 0.0001 (Wilcoxon rank sum test). **(f)** Two-photon shadow imaging in S1 L5 region of acute brain slices (P18). After the recovery, acute brain slices were perfused with Calcein solution (40 μM) in ACSF for 1 hour and then with Calcein solution in SeeDB-Live. See also Supplementary Video 6. **(g)** Inverse look-up table (LUT) images (left) and its magnified view (right) in the intermediate (100-115 μm stack) and deeper (185-200 μm stuck) regions. The LUT was optimized in each image. **(h)** Line plot showing the fluorescence along the dotted line in **(g)**. **(i)** The magnified view of the area with dense nerve fibers.

Previously, shadow imaging of organotypic brain slice cultures with super-resolution, confocal, and two-photon microscopy have been proposed for comprehensive structural imaging of the brain, including dense connectomics applications ^28–30^; in these techniques, the extracellular space of the brain slices is labeled with a dye solution (e.g. Calcein). The shadow images visualize the structure of all components in the tissue, allowing for comprehensive structural profiling. Previously, these techniques can only access the surface of the brain slices due to the light scattering. However, the surface of the brain slices (∼50 μm) is often mechanically damaged during slice preparation (Supplementary Fig. 5e), making it difficult to image “acute” brain slices that represent native *in vivo* structure. Using SeeDB-Live, the imaging depth possible for the shadow imaging was much improved with two-photon microscopy (Fig. 2f, Supplementary Video 6). Inverse look-up table (LUT) images demonstrated morphology of all the cells at higher signal-to-noise ratio under SeeDB-Live (Fig. 2g-i). Thus, SeeDB-Live facilitates comprehensive structural imaging of intact neuronal circuits.

### Electrophysiological properties of neurons cleared with SeeDB-Live

For comprehensive recording of neuronal activity with SeeDB-Live, it is important to ensure that neuronal functions remain intact. Using acute mouse brain slices (age, P14-18), we examined membrane properties of layer 5 pyramidal tract (L5PT) neurons using patch-clamp recording (Fig. 3a, Supplementary Fig. 6). The resting membrane potential of L5PT neurons was slightly lower under SeeDB-Live (−69.4 ± 2.9 mV; mean ± SD) than the control (−64.8 ± 2.2 mV) (Fig. 3b). This is most likely due to a slightly lower salt concentrations of SeeDB-Live (79.4% of ACSF) to maintain the osmolarity; indeed, this is in good agreement with the prediction based on the Goldman-Hodgkin-Katz voltage equation (assuming that intracellular ionic concentrations are not affected). We also found that the threshold membrane potential for the action potential is also slightly lower under SeeDB-Live (−48.1 ± 3.6 mV vs. −45.5 ± 4.2 mV) (Fig. 3b). As a result, the amplitude and frequency of action potentials upon current injection were comparable between control and SeeDB-Live (Fig. 3b, c). Thus, the firing properties of the neurons are largely preserved in SeeDB-Live.

**Figure 3.**
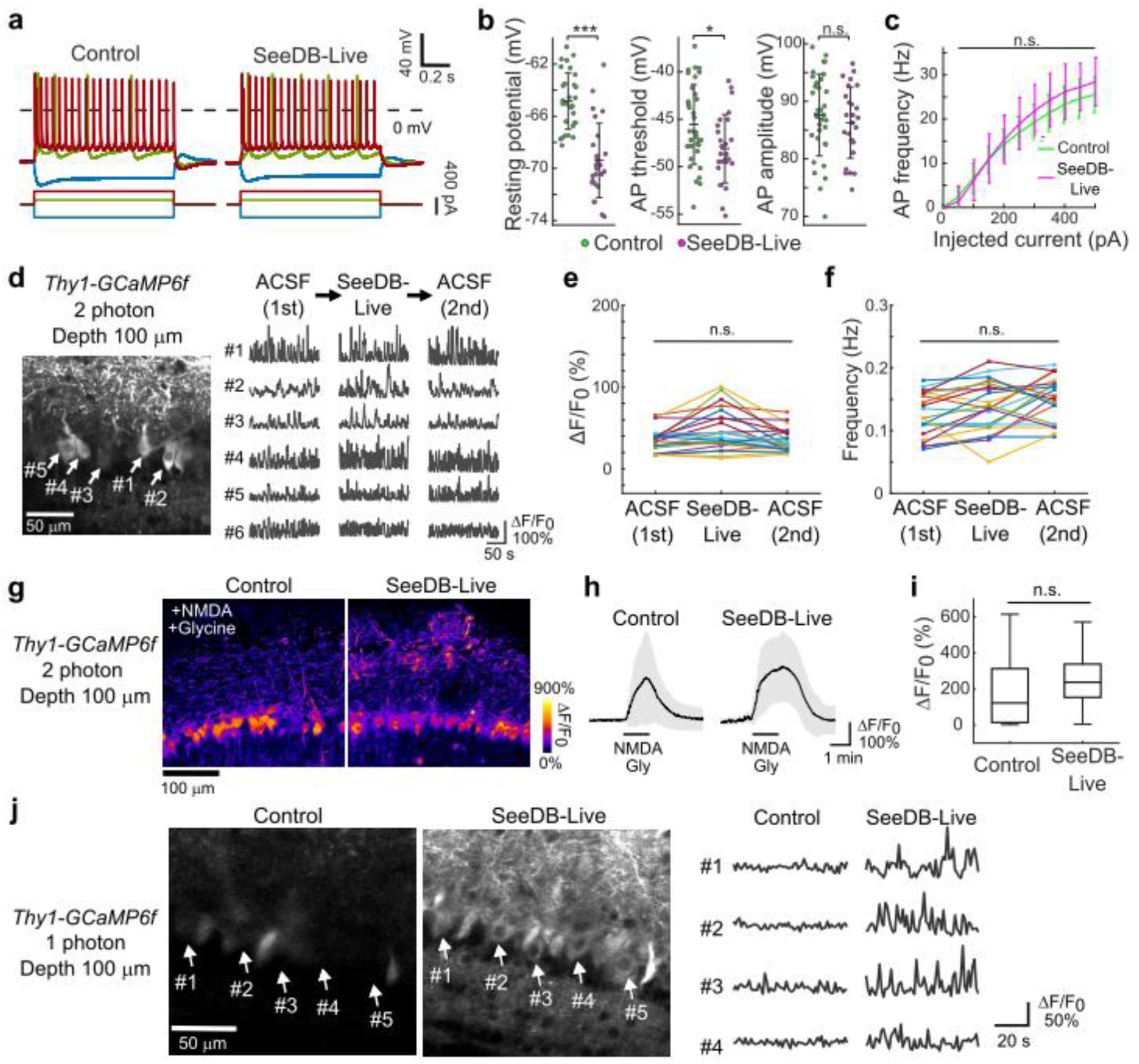
Electrophysiological recording and calcium imaging of neurons in acute brain slices. **(a)** Changes in membrane potentials in response to square current pulses of −300, +50, and +300 pA. Representative neurons are shown. The resting potentials were −66.5 and −68.2 mV for control and SeeDB-Live examples, respectively. **(b)** Population data for resting membrane potential, action potential (AP) threshold, and AP amplitude are shown. Data are means ± SD. \*\*\**p* < 0.001; \**p* < 0.05; n.s., not significant (*p* ≥ 0.05) (Wilcoxon rank sum test). n = 34 neurons for ACSF and n = 26 neurons for SeeDB-Live. **(c)** AP frequency was plotted against injected current amplitude. ACSF, n = 34. SeeDB-Live, n = 26. n.s., not-significant (Wilcoxon rank sum test corrected with Holm-Bonferroni correction). **(d)** GCaMP6f fluorescence images of mitral cells in acute brain slices (P11) imaged with two-photon microscopy. Spontaneous activity was imaged before (left), during (middle), and after (right) clearing with SeeDB-Live at a depth of 100 μm from the surface of the slice. Thy1-GCaMP6f mice were used. See also Supplementary Video 7. **(e, f)** Amplitude and frequency of spontaneous activity in the same set of mitral cells in ACSF and SeeDB-Live. n = 22 cells from 3 mice. n.s., non-significant (multiple comparisons with Bonferroni correction). **(g-i)** 100 μM NMDA and 40 μM Glycine were applied to the olfactory bulb slices (Thy1-GCaMP6f, age P15) for 1.5 min. n=70 and 47 cells for control and SeeDB-Live, respectively (3 slices). ΔF/F_0_ images **(g)**, time traces **(h)**, and response amplitudes **(i)** are shown. n.s., non-significant (student’s t-test). The slower decay of the response may be due to slower washout of NMDA/Gly under of SeeDB-Live. (**j**) Confocal images of acute olfactory bulb slices (Thy1-GCaMP6f mouse, age P13) before and after clearing with SeeDB-Live. SeeDB-Live improved the quality of images at a depth of 100 μm where most of the neurons are intact after slice preparation. Spontaneous activity was easily detected in SeeDB-Live, but poorly in the control. See also Supplementary Video 8.

### Calcium imaging of brain slices with SeeDB-Live

We next evaluated population-level properties of neurons using slice calcium imaging. We used olfactory bulb slices (P11-15), because mitral/tufted cells in the olfactory bulb show spontaneous activity in young animals ^31^. We used Thy1-GCaMP6f mice, in which mitral/tufted cells express GCaMP6f. We evaluated the physiological properties of mitral cells with two-photon microscopy at a depth of ∼100 μm. The frequencies of spontaneous activity were not significantly different between the control and SeeDB-Live when Ca^2+^/Mg^2+^ concentrations are optimized (Fig. 3d-f, Supplementary Video 7) (but see also Supplementary Fig. 2 for non-optimized conditions). In contrast, spontaneous activity was no longer maintained in the glycerol or iodixanol-containing ACSF (Supplementary Fig. 7). Evoked responses (to 100 μM NMDA and 40 μM glycine) of mitral cells were also comparable between control and SeeDB-Live (Fig. 3g-i). Spontaneous activity of mitral cells was clearly visible with one-photon confocal microscopy at a depth of 100 μm under SeeDB-Live, but not with control ACSF (Fig. 3j, Supplementary Video 8). It should be noted that the superficial ∼50 μm of acute brain slices is typically damaged during sample preparation and intact neuronal activity is only visible in deeper areas, where only two-photon microscopy can access under the normal ACSF. Thus, SeeDB-Live enables calcium imaging of healthy neuronal activity in acute brain slices using conventional confocal microscopy, without using expensive two-photon microscopy systems.

### Optical clearing of cerebral cortex *in vivo*

SeeDB-Live is based on index matching with a membrane-impermeable molecule, BSA. Therefore, the performance of clearing is limited by the accessibility of BSA to the tissues. To clear the mouse brain *in vivo* in live animals, we performed a craniotomy at the primary somatosensory cortex (S1) and removed the dura mater (durotomy). We then exposed the brain surface to SeeDB-Live/ACSF, allowing BSA to permeate into the cerebrospinal fluid of the brain (Fig. 4a). We confirmed that fluorescently tagged BSA was infused to a depth of

**Figure 4.**
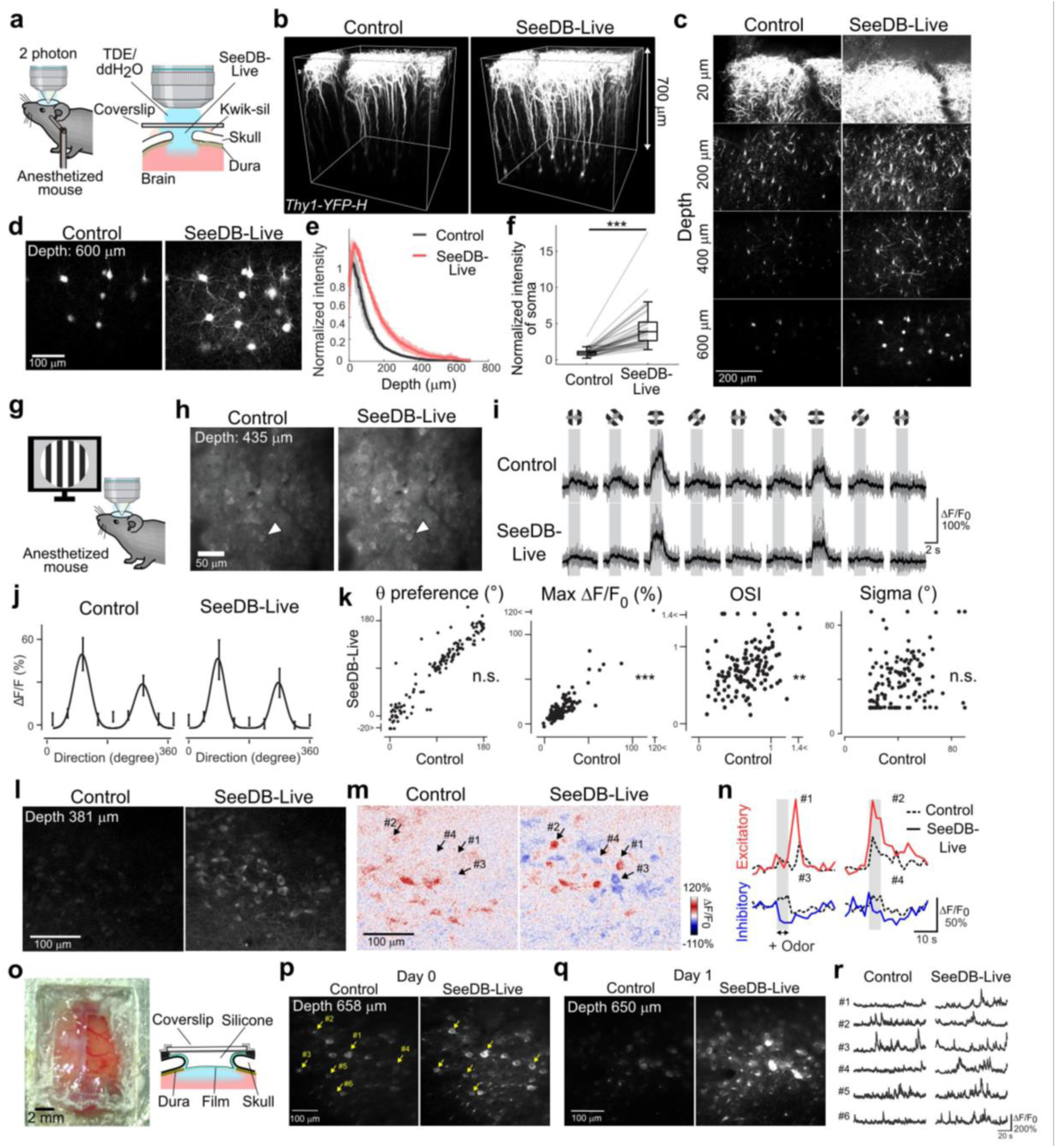
Improved imaging of intact brain functions with SeeDB-Live *in vivo*. **(a)** Schematic diagram of surgery and clearing of the mouse cortex with SeeDB-Live (refractive index 1.366, 300 mOsm/kg in ACSF-HEPES). Craniotomy (5 mm diameter) was made on S1, and dura was removed. 100-200 μL of SeeDB-Live was held in the imaging chamber. TDE/ddH_2_O (refractive index 1.366) was used for the immersion. Supplementary Fig. 8 shows the diffusion of fluorescently labeled BSA into the cortex. **(b)** Primary somatosensory cortex (S1) of Thy1-EYFP-H mouse (6-month-old) was imaged before and after clearing with SeeDB-Live (1 hour after clearing) with two-photon microscopy. Layer 5 pyramidal tract (L5PT) neurons are labeled with EYFP in Thy1-YFP-H mice. **(c)** *x-y* images at different depths. See also Supplementary Video 9. **(d)** Basal dendrites of L5PT neurons were clearly visualized after SeeDB-Live treatment. **(e)** Fluorescence intensity at different depths. The mean fluorescence signals were determined for each depth. **(f)** Fluorescence intensities of L5PT neurons in S1 before and after clearing with SeeDB-Live. Somatic signals from the same sets of neurons were compared. \*\*\**p* < 0.0001 (paired Student’s t-test). **(g)** Two-photon calcium imaging of L2/3 neurons expressing jGCaMP8m in the primary visual cortex (V1) before and after clearing with SeeDB-Live. See also Supplementary Fig. 9a-e for Cal-520 data. **(h)** Basal fluorescence of jGCaMP8m without visual stimulation. L2/3 neurons at a depth of 435 μm. **(i, j)** Responses of a representative L2/3 neuron (indicated by arrowheads in **h**) to visual grating stimuli before and after clearing with SeeDB-Live. **(k)** Preferred orientation, maximum responses (ΔF/F_0_), orientation selective index (OSI), and tuning width (Sigma) for the same set of L2/3 neurons (136 neurons in total) before (*x*-axis) and after (*y*-axis) clearing with SeeDB-Live. The comparison was performed as described previously ^63^. Orientation tuning was not affected by clearing with SeeDB-Live. \*\*\**p* < 0.001; \*\**p* < 0.01; n.s., non-significant (Wilcoxon signed-rank test). **(l)** x-y images of mitral cells in the olfactory bulb of Thy1-GCaMp6f mouse (4-month-old) was imaged before and after clearing with SeeDB-Live (1 hour after clearing) with two-photon microscopy. The depth was 381 μm. See also Supplementary Video 10. **(m)** Odor responses of mitral cells upon 1% valeraldehyde. Excitatory and inhibitory responses were clearly observed after SeeDB-Live treatment. Arrows indicate the same sets of neurons. **(n)** Representative excitatory/inhibitory responses of mitral cells indicated in **(m)** before and after SeeDB-Live treatment. **(o)** A photo (left) and schematic diagram (right) of a large cranial window with a plastic film. After clearing with SeeDB-Live for 1 h, the brain surface was covered with a plastic film. ddH_2_O was used for the immersion. **(p)** Basal fluorescence (temporal median) of RCaMP3-expressing L5 neurons before and after SeeDB-Live treatment at Day 0. The depth was 658 μm. **(q)** Basal fluorescence of RCaMP3-expressing L5 neurons at Day 1. The depth was 650 μm. **(r)** Representative Ca^2+^ responses of L5 neurons indicated in **(p)** during repeated whisker stimuli.

∼800 μm from the surface of the cerebral cortex (Supplementary Fig. 8). We used a transgenic line, Thy-YFP-H, in which layer 5 pyramidal tract (L5PT) neurons are labeled with EYFP. Under two-photon microscopy, overall brightness of EYFP signals was increased after incubation with SeeDB-Live for 1 hour (Fig. 4b, c, Supplementary Video 9). Their basal dendrites were better visualized with SeeDB-Live (Fig. 4d). The brightness of L5PT somata, located at a depth of 600-800 μm, was increased ∼4-fold by SeeDB-Live treatment (Fig. 4e, f). Thus, SeeDB-Live is a powerful tool for *in vivo* imaging of live neurons in the brain.

### *In vivo* calcium imaging of neurons with SeeDB-Live

Next, we investigated whether sensory responses are preserved after SeeDB-Live treatment *in vivo*. We performed a durotomy in the primary visual cortex (V1) and compared the visual responses of layer 2/3 neurons before and after SeeDB-Live treatment (Fig. 4g), assuming that layer 2/3 is fully infused with SeeDB-Live (Supplementary Fig. 8). Using a calcium indicator, jGCaMP8m and Cal-520, we recorded the calcium responses of the same sets of neurons to visual grating stimuli of different orientations (Fig. 4h-k, Supplementary Fig. 9a-e). We found no obvious changes in the preferred orientation, response amplitude (ΔF/F_0_), orientation selective index, and tuning width of layer 2/3 neurons (Fig. 4i-k). Thus, SeeDB-Live preserves physiological sensory responses.

In the olfactory bulb, we were able to better visualize odor responses in mitral cell somata located at a depth of ∼400 μm using GCaMP6f (Fig. 4l). Due to the improved brightness, inhibitory responses, represented by a reduction in basal GCaMP6f fluorescence, were better detected using SeeDB-Live (Fig. 4m, n, Supplementary Video 10).

Optical clearing with SeeDB-Live is transient *in vivo* as BSA is gradually washed out. To perform chronic calcium imaging with SeeDB-Live in awake animals, we used easily removable plastic films for the cranial window. A large cranial window (6 x 3 mm^2^) was made in one hemisphere and SeeDB-Live was applied for 1 hour. Then, a polyvinylidene chloride (PVDC) wrapping film was attached onto the window, which was then stabilized with silicone sealant and a coverslip ^32^. We obtained stable responses of RCaMP3 ^33^ from layer 5 in S1 (650 μm depth). Since the window is easily detached, we could repeat the same procedures on consecutive days without compromising the quality of the window (Fig. 4o-r).

### Widefield voltage imaging of acute brain slices with SeeDB-Live

Calcium is an indirect measure of neuronal activity. Genetically encoded voltage indicators (GEVIs) with high signal-to-noise ratios have been developed in recent years. To image fast voltage changes, high-speed widefield imaging is advantageous over point-scanning two-photon microscopy. Here we cleared acute olfactory bulb slices with SeeDB-Live and imaged calcium and voltage signals using GCaMP6f and a fast and sensitive chemigenetic voltage indicator, Voltron2, sparsely introduced to mitral/tufted cells by *in utero* electroporation (Fig. 5a) ^34^; Voltron2 was visualized with JF_549_-HaloTag ligand applied to the medium (Voltron2_549_). After the clearing with SeeDB-Live,, the epifluorescence signals of Voltron2_549_ were clearly visualized at a depth of > 150 μm (Fig. 5b). Using a high-speed CMOS camera (2 kHz), we recorded voltage changes in different compartments of mitral cell dendrites: We could visualize the back-propagation of action potentials from somata to dendritic tips in single-shot imaging, without averaging (Fig. 5c-f, Supplementary Video 11). We observed a ∼1.5 ms delay in responses at the tip of the primary dendrites (Fig. 5g). Thus, SeeDB-Live will be a powerful tool for studying subcellular dynamics of voltage signals in acute brain slices.

**Figure 5.**
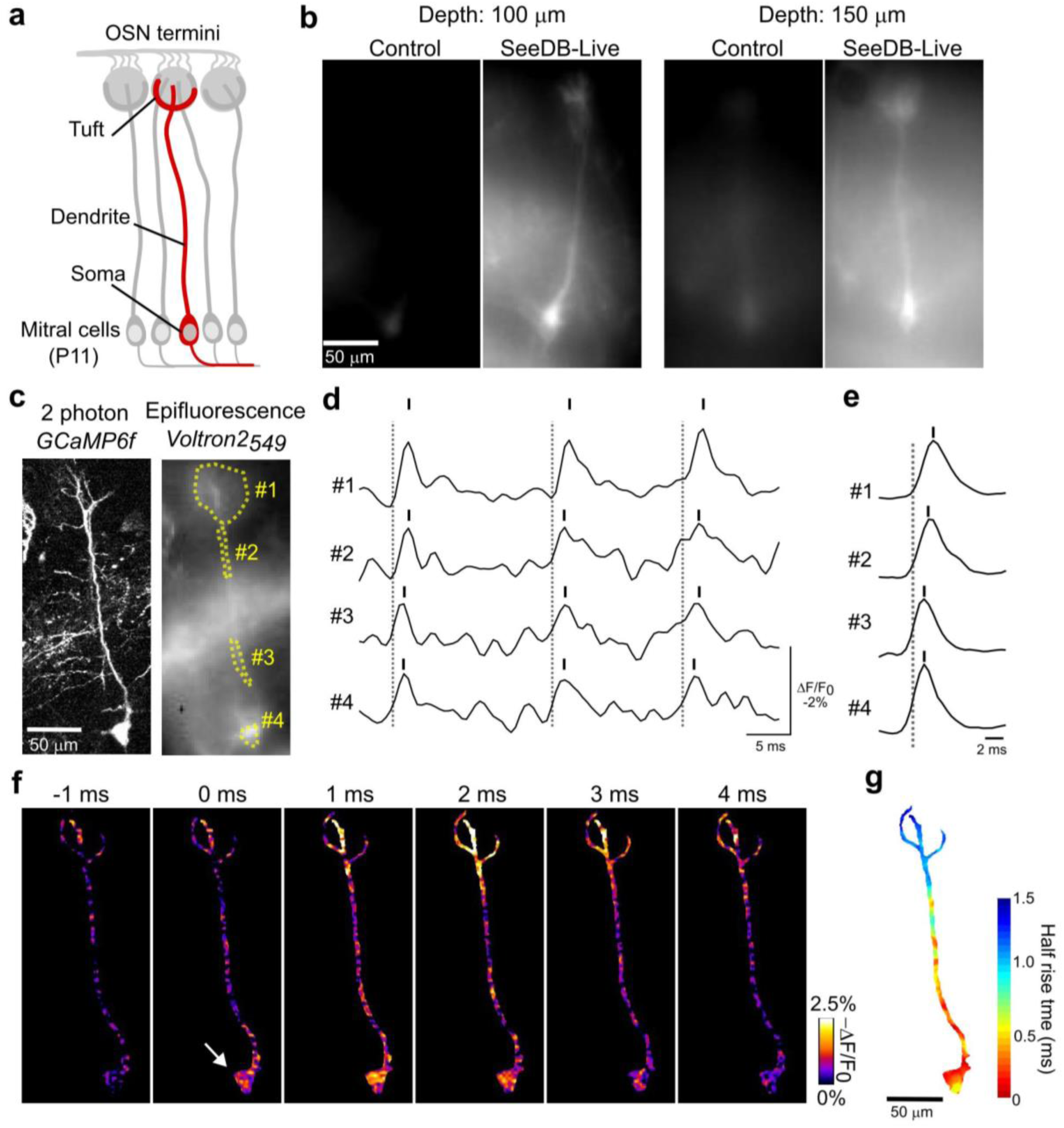
Widefield imaging of subcellular voltage changes of neurons in acute brain slices. **(a,b)** Voltage imaging of a mitral cells using epifluorescence microscopy **(a)**. Mitral cells labeled with Voltron2_549_ at different depths before and after SeeDB-Live treatment (acquired with a high-speed CMOS camera) **(b)**. Voltron2 (labeled with Janelia Fluor HaloTag Ligand549) were introduced to mitral cells by *in utero* electroporation, and olfactory bulb slices were imaged at P11. **(c)** Two-photon image identified a labeled mitral cell (left). Epifluorescence of Voltron2_549_ and ROIs are shown on the right. Cell bodies, shafts of a primary dendrite, and tufted structure within a glomerulus were analyzed. **(d,e)** Time traces of backpropagating action potentials in each ROI indicated in **(c)**. Traces from single shot images **(d)** and averaged images from 65 events **(e)** are shown. Ticks indicate the peaks of detected action potentials. Dotted lines indicate the half rise time of the peaks from ROI#4. **(f)** Backpropagation of action potentials were imaged at 2 kHz (single shot images). Representative −ΔF/F_0_ images are shown. ROI was manually cropped based on 2-photon and widefield images. See also Supplementary Video 11. **(g)** Spatiotemporal pattern of backpropagating action potentials, averaged from 65 events. The half-rise timing of action potentials at soma was defined as 0 ms. ROI was manually cropped based on 2-photon and widefield images. Median filtering (4×4 pixels) was applied to the images.

### Widefield voltage imaging of backpropagating action potentials *in vivo*

Previously, it has been challenging to image GEVI signals at deeper part of the brain using widefield imaging *in vivo* ^34–36^. Here we expressed Voltron2-ST specifically in layer 2/3 (L2/3) pyramidal neurons in the primary somatosensory cortex (S1) using *in utero* electroporation, visualized with JF_549_-HaloTag ligand. After the clearing with SeeDB-Live, L2/3 neurons located at a depth of 120-150 μm were better visualized, allowing for reliable detection of spontaneous action potentials in their somata (Supplementary Fig. 9f-h). We also detected backpropagating action potentials from dendrites of L2/3 neurons in awake mice (Supplementary Fig. 9i-k).

Next, we used SeeDB-Live to perform voltage imaging from a population of neurons *in vivo*. In the olfactory bulb, the odor information is encoded not only by the spike frequencies, but also by the timing ^37^. However, in previous electrophysiological studies, it has been difficult to relate the recorded spike data to the anatomical information. Widefield voltage imaging has been performed in the olfactory bulb, but never at the single neuron resolution ^38, 39^.

We expressed Voltron2 specifically and sparsely in mitral/tufted cells using the Pcdh21-Cre driver and Cre-dependent AAV vector, labeled with JF_549_-HaloTag ligand (Voltron2_549_). Voltron2_549_ signals were found not only in somata, but also in dendrites (Fig. 6a-c). As a result, we were able to detect backpropagating action potentials from ∼140 neurites (ROIs) at a depth of ∼90 μm using SeeDB-Live and widefield imaging (Fig. 6d). In this imaging setup, the theoretical focal depth was ∼1.36 μm (Fig. 6a). Since the fluorescence changes were up to 2-3% ΔF/F_0_ (much smaller than calcium imaging), it is unlikely that out-of-focus signals interfere or contaminate the spike detection in each of the ROIs.

**Figure 6.**
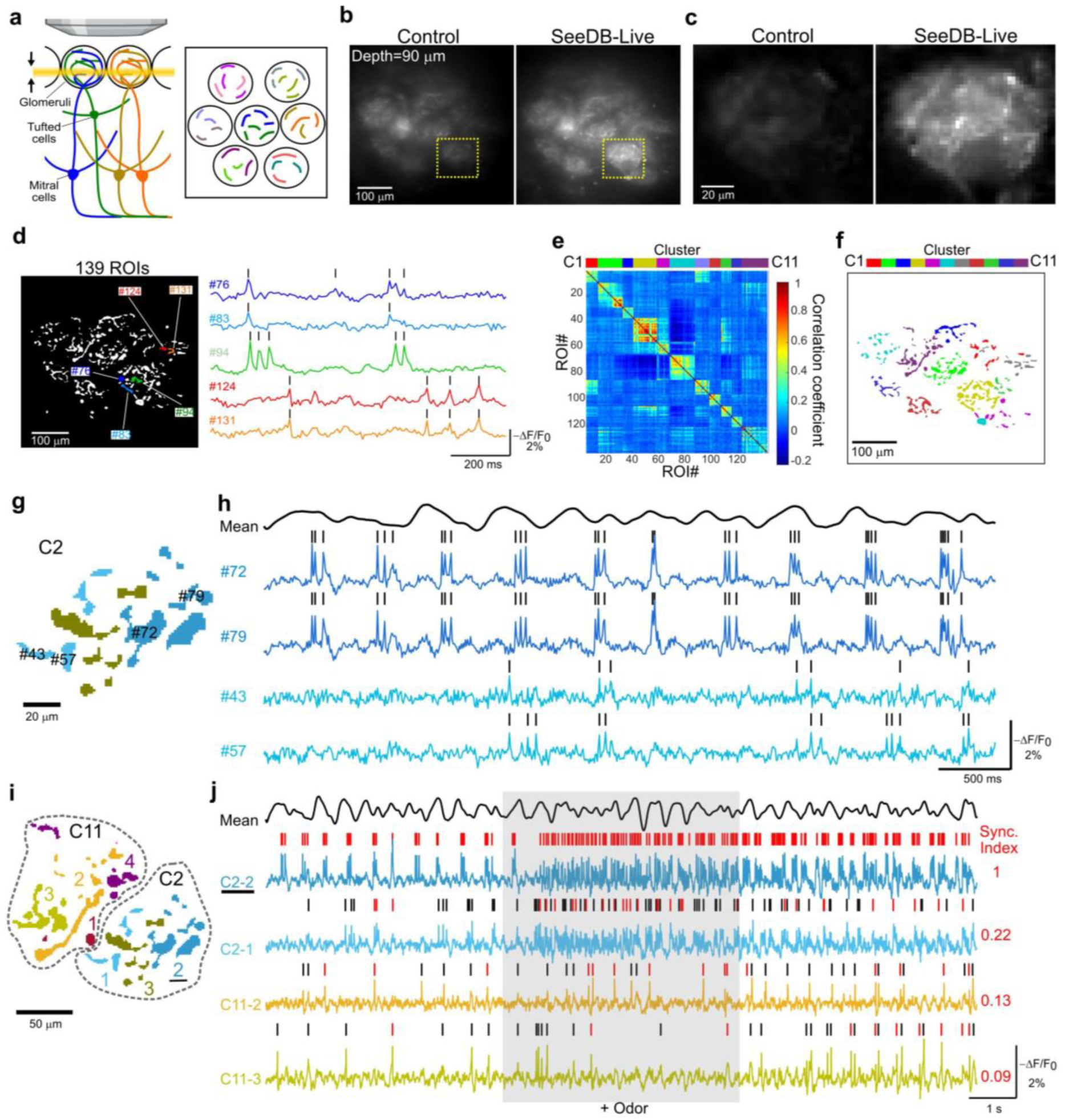
Widefield population recording of voltage changes in the mouse brain *in vivo*. **(a)** Schematic diagram of widefield voltage imaging in the mouse olfactory bulb *in vivo*. Mitral/tufted cells were sparsely labeled with Voltron2_549_. We imaged a deeper part of the glomerular layer (90 μm), where mitral/tufted cells form dendritic branches within the glomerulus. A focal plane is shown as a yellow line (arrows). We used a 25x objective (NA 1.05) with a short focal depth (1.36 μm) to minimize out-of-focus signals. It is unlikely that scattered out-of-focus signals contaminate detection of spikes at the focal plane, since the response amplitude is only up to 2-3 %, contrary to calcium imaging. **(b)** Epifluorescence images of dendrites (and some somata) of mitral/tufted cells labeled with Voltron2_549_ in an anethetized mouse (4-month-old) before and after clearing with SeeDB-Live (1 hour after clearing). F_0_ images at a depth of 90 μm. **(c)** Boxed areas in **b** are shown. **(d)** ROIs were semi-automatically detected using machine learning-based image segmentation (ilastik) of the F_0_ image. See Supplementary Video 12 for Voltron2_549_ −ΔF/F_0_ images in ROIs. Representative traces from highlighted ROIs (dendrites) are shown on the right. Voltron2_549_ −ΔF/F_0_ is shown. Ticks indicate the detected action potentials. Note that subthreshold activities were also correlated between ROIs within the same glomerulus. **(e)** Cross-correlation matrix for voltage traces from ROIs. ROIs were clustered using k-means clustering. The cluster number k was defined based on the number of glomeruli. **(f)** Spatial distribution of ROIs in each cluster. Each color represents a different cluster. **(g,h)** Subclusters based on spike synchronicity between ROIs within a glomerulus (Cluster 2) **(g)** and representative traces from indicated ROIs **(h)**. The black trace on the top (Mean) shows the averaged −ΔF/F_0_ of all glomeruli indicating sniff-coupled theta wave in the olfactory bulb. **(i,j)** Comparison of synchronicity between the subclusters in the same or different glomeruli (Cluster 2 and 11) **(i)** and representative traces of the indicated subclusters **(j)**. The red ticks indicate the synchronous events that coincided with spikes in the C2-2 subcluster. The synchronicity index indicates the proportion of synchronous events normalized by the spike frequency. Odor (1% amyl acetate) was delivered to the mice during the gray shaded period.

Correlation of the voltage traces across ROIs revealed ∼11 discrete clusters (Fig. 6e). The ROIs within each cluster (11 clusters by k-means) were also spatially clustered (Fig. 6f), demonstrating that neurites within the same glomerulus have similar voltage dynamics (including subthreshold changes) (Fig. 6d). This makes sense because dendrites connecting to the same glomerulus receive similar synaptic inputs and are electrically coupled within the glomerulus ^40^. When we looked at individual ROIs within the same glomerulus, some pairs, but not all, demonstrated highly correlated backpropagating action potentials, suggesting that these dendritic branches originated from the same neuron (Fig. 6g, h). We therefore grouped ROIs with highly synchronized backpropagating action potentials into a subcluster (see **Methods**). In this way, we obtained 21 subclusters that most likely represent different neurons (Supplementary Fig. 10). Subclusters that belong to the same glomerulus (representing “sister” mitral/tufted cells connecting to the same glomerulus) tend to show more synchronized events than those in different glomeruli (Fig. 6i, j) ^40^. We also observed odor-evoked phase shifts in action potentials relative to the sniff-coupled theta oscillations (Supplementary Fig. 10c), consistent with previous studies ^41, 42^. Thus, widefield imaging of dendrites combined with SeeDB-Live provides a powerful approach for studying population voltage dynamics *in vivo*.

## Discussion

In this study, we found that the addition of spherical polymers, especially BSA, to the extracellular media is effective in making live cells transparent without affecting osmolarity. We demonstrated the power of our live tissue clearing approach in confocal, widefield, and two-photon microscopy for spheroids, organoids, acute brain slices, and mouse brain *in vivo*. With SeeDB-Live, fluorescence brightness was increased, and imaging depth was improved. Combined with wider field-of-view two-photon microscopy ^43–46^, targeted one-photon imaging approaches ^35, 36^, and red-shifted indicators ^33^ , SeeDB-Live expands the imaging scale for tissue and organ-scale biological phenomena both *ex vivo* and *in vivo*.

In this study, we demonstrated that SeeDB-Live is particularly useful for widefield voltage imaging. Previously, large-scale imaging of voltage changes has been difficult due to the slow scanning speed of two-photon microscopy and light scattering with one photon microscopy. We have shown that SeeDB-Live allows for reliable detection of action potentials with widefield one-photon imaging *in vivo*. Moreover, we demonstrate that we can record backpropagating action potentials reliably from dendrites. Widefied imaging of dendritic voltage changes with SeeDB-Live would be a powerful strategy for studying subcellular and/or population-scale voltage dynamics both *ex vivo* and *in vivo*.

We have demonstrated the utility of SeeDB-Live for functional assays of organoids. For the improved culture of organoids, microfluidic culture systems may be useful to improve circulation and/or to exchange the medium ^26^. For the *in vivo* applications, accessibility of BSA to the target tissues may be the major issue. For the chronic *in vivo* imaging of the mouse brain, we demonstrate the utility of easily removable cranial windows ^32^. In the future, BSA-permeable membrane may be more useful. Alternatively, infusion into the cerebrospinal fluid system may be useful for efficient permeation of SeeDB-Live for more extensive clearing of the entire brain in the future studies ^47^. Other organs may be more difficult to clear, and future *in vivo* applications would require strategies to overcome the accessibility issue.

The improved transparency with SeeDB-Live also expands the modality of the imaging methods. With SeeDB-Live, we can now use affordable confocal microscopy for deep imaging, enabling high-resolution multicolor imaging. The combination of fluorescence imaging and photostimulation will be easier with one-photon and SeeDB-Live than with a multi-photon setup. As optical aberration is minimized, SeeDB-Live should also be very useful for super-resolution imaging of large volume in live tissues ^9, 48^. While we have demonstrated shadow imaging with two-photon microscopy, STED microscopy in combination with SeeDB-Live may enable saturated connectomics in acute brain slices, rather than in cultured brain slices^29, 30^. For the best performance, it will be important to design objective lenses optimized for SeeDB-Live (refractive index ∼1.366). Together with ongoing efforts to develop microscopy techniques, our live tissue cleaning approach facilitates our understanding of the tissue- and organ-scale dynamics of biological phenomena.

## Methods

### Mice

All animal experiments were approved by the Institutional Animal Care and Use Committee (IACUC) of Kyushu University, Kagoshima University, and Yamanashi University. A *Thy1-GCaMP6f* Tg (line GP5.11) (JAX #024339) ^49^, *Thy1-YFP-H* (JAX #003782) ^50^ and *Pcdh21-Cre* (RIKEN BRC, RBRC02189) ^51^ have been described previously. ICR, C57BL6J and C57BL/6N mice were purchased from Japan SLC. Both males and females were used for our experiments. Mice were kept under a consistent 12-hour light/12-hour dark cycle (lights on at 8 a.m., and off at 8 p.m.).

### Plasmids

To construct pCAG-GCaMP6f, GCaMP6f gene was PCR amplified from pGP-CMV-GCaMP6f (Addgene #40755) with Q5 High-Fidelity 2X Master Mix (M0492S, NEB). The cDNA was flanked by *EcoR*I and *Not*I sites. The GCaMP6f cDNA was subcloned into pCAG vector with Ligation Kit (6023, Takara). To make pAAV-CAG-jGCaMP8f-WPRE, jGCaMP8f gene was amplified from pGP-AAV-syn-jGCaMP8f-WPRE (Addgene #162376) with Q5 High-Fidelity 2X Master Mix. The cDNA contained an extra 20-30 bp overlap regions with pAAV-CAG-tdTomato (Addgene #59462). The tdTomato was removed from pAAV-CAG-tdTomato by digestion with *Kpn*I and *Hind*III, and jGCaMP8f cDNA with the extra sequence was subcloned into the vector with NEBuilder HiFi DNA Assembly (E2621S, NEB). Plasmids used in this study will be deposited to Addgene (https://www.addgene.org/Takeshi_Imai/).

### Preparation of SeeDB-Live

Crystallized bovine serum albumin (BSA) from bioWORLD (#22070004) was used for SeeDB-Live. BSA was dissolved in culture medium or ddH_2_O with gentle shaking. Note that stirring tends to produce precipitation. Additional CaCl_2_ and MgCl_2_ were supplemented after BSA was fully dissolved in the medium (except for Fig. 1a-k and Supplementary Fig. 1): The ratio of the additional divalent cations was the same as in the original medium, and the total amount was +3 mM. As for the medium for intestinal culture (IntestiCult Organoid Growth Medium), the ratio of the additional CaCl_2_ and MgCl_2_ was 2 : 1 (Fig. 1o-q, Supplementary Fig. 4e-f). The osmolarity of the BSA solution was measured with a vapor pressure osmometer (VAPRO 5600, Xylem ELITech). The final concentration of the medium/saline was adjusted to 80-90% of the original medium/saline so that the osmolality of the BSA-containing medium/saline would be the same as that of the original medium/saline. The refractive index was measured with an Abbe refractometer (ER-2S, Erma) with a white LED light source. The refractive index of the clearing media was adjusted to 1.363-1.366. To oxygenate the medium, 95% O_2_/ 5% CO_2_ gas was filled in the bottle containing the medium for 1-2 hours before the experiments. Saturation of O_2_ was checked with an O_2_ sensor (#9521, Horiba). Bubbling is not recommended because it produces a lot of foams, and BSA may be denatured. Notably, different lots of commercialized BSA demonstrated different levels of autofluorescence and precipitation. For quality check, we first measured the autofluorescence of 15% (w/v) BSA/ddH_2_O solutions and selected the lots showing the lower autofluorescence at 488 nm excitation using a fluorescence spectrophotometer (F2700, Hitachi High-Tech). We then selected lots showing no precipitation at 15% (w/v) BSA/ACSF.

We tested following BSA products: BSA#1, BSA crystal (#22070004, bioWORLD); BSA#2, BSA crystal (#012-15093, Fujifilm); BSA#3, BSA Low Salt (# 015-15125, Fujifilm); BSA#4, BSA pH 5.2 (# 017-21273, Fujifilm); BSA#5, BSA pH 7.0 (# 019-27051, Fujifilm); BSA#6, BSA Globulin Free (# 016-15111, Fujifilm); BSA#7, BSA Protease Free (# 019-28391, Fujifilm).

We tested the following chemicals during the screening process: Glycerol (#17018-25, Nacalai), DMSO (#043-07216, Fujifilm), Propylene glycol (#164-04996, Fujifilm), iodixanol (VISIPAQUE 320 INJECTION 50 mL, GE health care), iodixanol (#D1556-250ML, Optiprep), Iotrolan (Isovist Inj. 300, Bayer Pharma Japan), Iopamidol (#OYPALOMIN, FujiPharma), Iopromide (#Iopromide 370 Injection [FRI], Fujifilm), Iohexol (OMNIPAQUE350 INJECTION, GE healthcare Pharma), Ioxilan (Imagenil350 inj., Guerbet Japan), Ioversol (Optiray350 Inj., Mallinckrodt), Ioxagilic acid (#Hexabrix320 inj., Guerbet Japan), Iomeprol (Iomeron400 Bracco-Eisai), Ficoll 70 (#17031050, Cytiva), Ficoll400 (#17030010, Cytiva), HBCD (#307-84601, Glico), PVP (#P0471, TCI), Sucrose (#193-00025, Fujifilm), Polydextrose (#polydex300, Nichiga), Partially Hydrolyzed Guar Gum (PHGG, # 2021092403, Nichiga), Methyl-β-cyclodextrin (Mβ-CD, #M1356, TCI), Agave inulin (#agabe500, Nichiga), PEG8000 (#V3011, Promega), RM+Stevia (#dex-5-500m, Nichiga), Resistant maltodextrin from corn (RM1, #MK-H108-6T6I, Nichiga), Resistant maltodextrin from wheat (RM2, #dekisutorin-komugi-400, Nichiga), Inulin (#inurinn500, Nichiga), Isomaltodextrin (Fibryxa, Hayashibara), Reduced resistant maltodextrin (RRM, #kg-nandeki-400, Nichiga) , Stevia (#sutebiasw5-150m, Nichiga).

See Supplementary Table 1 (Excel file) for more detailed composition of SeeDB-Live and other clearing media.

### Transmittance measurement in cell suspension

HeLa/Fucci2 cells (RCB2867, Riken BRC) were cultured in Dulbecco’s Modified Eagle Medium (DMEM high glucose) (#043-300085, Fujifilm) supplemented with 1% Penicillin-Streptomycin Solution and 10% fetal bovine serum (FBS) at 37°C, 5% CO_2_. After trypsinization, cells were collected at 2 × 10^5^ cells/tube. After centrifugation, the medium was replaced with 50 μL of index-adjusted PBS. Transmittance of the cell suspension in 400-1100 nm was measured with a ratio beam spectrophotometer (U-5100, Hitachi High-Tech).

### Ca^2+^ measurement in cultured cells

HEK293T cells (AAVpro 293T, #632273, Takara) were cultured in DMEM (high glucose) supplemented with 1% Penicillin-Streptomycin Solution and 10% FBS at 37°C, 5% CO_2_. Cells seeded in 35 mm glass bottom dishes (60% confluent) were transfected with pGP-CMV-GCaMP6f (Addgene#40755) using PEI Max (Polysciences, Inc., #24765-1). Twenty-four hours after transfection, the medium was replaced with an index-adjusted medium (refractive index 1.365). The osmolality of the medium was not adjusted to isotonicity. Two hours after the medium exchange, cells were imaged with an inverted microscope (DMI600B, Leica) equipped with a 10x NA0.4 dry objective lens. 50 μM ATP (final concentration) was added to the medium during imaging. The maximum values after stimulation were used for the data analysis. For the calcium measurement with a plate reader (TriStar LB941, Berthold), GCaMP6f-expressing HEK293T cells were transferred to a 96-well plate at 4 × 10^5^ cells/mL/well. 50 μM ATP was added to the medium during the time-series measurement. Mean values during 1-10 sec after stimulation were analyzed.

### Viscosity measurement

To measure the viscosity of the solutions, we measured the time taken for 20 mL of solutions to flow out from a 50 mL syringe (TERMO, #SS-50ESZ) with an internal tip diameter of ∼2 mm tip. The point at which the solution flow was stopped was considered as the end of the flow. The viscosity of sucrose solutions at room temperature (21°C) ^52^ was used as the standard. The plots were fitted by single exponential fitting. The viscosity of the solutions was calculated based on the calibration curve.

### Cell growth measurement

HeLa/Fucci2 cells were seeded on a clear bottom 384-well plate at 700 cells/well. The cells were cultured in DMEM (high glucose), without phenol red, and glutamine (#040-30095, Fujifilm) supplemented with 1% Penicillin-Streptomycin Solution, 10% FBS, and 1% glutamine at 37°C, 5% CO_2_. Twenty-four hours after seeding, the medium was replaced with an index-adjusted medium (refractive index 1.363, 310-320 mOsm/kg). Fucci2 fluorescence (mVenus and mCherry) was imaged with an inverted fluorescence microscope (DMI600B, Leica) equipped with a 5x NA 0.1 dry objective lens. Cells were counted based on the nucleus images of Fucci2 fluorescence with Image J software (https://imagej.net/ij/). First, the green and red channels were summed. The speckle noise was removed with “Despeckle”. The overlapped nuclei were separated with “Watershed”. The intensity threshold was determined manually. Finally, the cell number in each well was counted with “Analyze particles”. For manual counting, the cells were seeded on 35 mm dishes at 1 × 10^5^ cells/dish. Twenty-four hours after plating, the medium was replaced with an index-adjusted medium (refractive index 1.363, 320 mOsm/kg). After trypsinization, the cell number was counted with a hemocytometer.

For spheroid formation, HeLa/Fucci2 cells were seeded on an ultra-low attachment 96 well plate (#7007, Corning) at 1000 cells/well. 24-48 hours after plating, the spheroid was incubated in an index-adjusted medium (SeeDB-Live, refractive index 1.366, 320 mOsm/kg) for four hours per day or for all the time. The half volume of the culture medium was replaced with the fresh one every day for the organoid culture in Fig. 1o,p. For manual counting, the cells were incubated in a mixture of 50 μL DMEM and 200 μL Trypsin-EDTA for 30 min. The suspension was then centrifuged at 1000 rpm for 5 min. The pellet was resuspended with the culture medium for cell count with a hemocytometer.

### Spheroid imaging

HeLa/Fucci2 spheroid was incubated in SeeDB-Live (refractive index 1.366, 320 mOsm/kg) for one hour. Then, the spheroid was mounted on a glass slide and sealed with a 1 mm thickness silicone rubber spacer (Togawa rubber) and a coverslip (Matsunami). Imaging was performed with FV1000MPE microscope (Olympus/Evident) with Fluoview FV10-ASW software (Olympus/Evident, RRID:SCR_014215) and a 25x NA 1.05 objective lens (Olympus/Evident, XLPLN25XWMP). Immersion was water or TDE/ddH_2_O (refractive index 1.366) for control or SeeDB-Live samples, respectively. The correction collar was turned to the appropriate position. For the confocal imaging, 473 nm and 569 nm lasers were used to excite mVenus and mCherry, respectively. For the two-photon imaging, a femtosecond laser (Insight DeepSee, SpectraPhysics) was tuned to 920 nm for mVenus excitation. 1040 nm laser was used for mCherry excitation.

For cell detection, flat areas of the images were cropped. The green and red channels were merged to make reference images. A median filter was applied (2 × 2 pixels). Regions of interest (ROIs) for each cell in a spheroid were created with Cellpose ^53, 54^. Cell numbers and the fluorescence intensity were calculated based on the ROIs using MATLAB software (MathWorks). 3D-rendered images were made by ImarisViewer (Oxford Instruments).

### Imaging of intestinal organoids

Intestinal organoids were created following the manufacturer’s protocol (VERITAS). Briefly, ePET-Cre; Ai162 mice were sacrificed with an overdose i.p. injection of pentobarbital (100-150mg/kg). A small intestine was taken out and cut to expose the lumen side. The lumen was gently washed with cold PBS (-) several times. The small intestine was cut into 2 mm pieces in 10 mL of cold PBS (-). After pipetting 3 times, the supernatant was replaced with new cold PBS (-). This procedure was repeated >15 times until the supernatant became clear. The supernatant was replaced with 25 mL of Gentle Cell Dissociation Reagent (ST-100-0485, STEMCELL Technologies). The pieces were gently shaken for 15 min at room temperature. The supernatant was replaced with 10 mL of cold 0.1% BSA/PBS. After pipetting 3 times, the suspension was passed through a 70 µm Cell Strainer (352350, Corning). This step was repeated to obtain the fraction containing more crypts. After centrifugation at 300 g for 5 minutes at 4°C, the supernatant was replaced with 10 mL of cold 0.1% BSA/PBS. After centrifugation at 200 g for 3 minutes at 4°C, the supernatant was replaced with 10 mL of DMEM/F-12 (#11039-021, ThermoFisher). After centrifugation at 200 g, for 5 min at 4°C, the supernatant was replaced with 150 µL of IntestiCult Organoid Growth Medium (ST-06005, STEMCELL Technologies). 150 µL of Matrigel (#356237, Corning) was added to the suspension. After pipetting 10 times, 50 µL of the mixture was mounted on the well of a 24-well plate. The plate was incubated for 10 min at 37°C to gelatinize the Matrigel. 750 µL of IntestiCult Organoid Growth Medium was added to the wells carefully.

For imaging, the organoids were dissociated from a gel by pipetting with Gentle Cell Dissociation Reagent and transferred to a 15 mL tube. The tube was gently shaken for 10 min at room temperature. After centrifugation at 300 g for 5 minutes at 4°C, the supernatant was replaced with 10 mL of cold DMEM-F12. After centrifugation at 300 g, for 5 min at 4°C, the supernatant was replaced with 150 µL of IntestiCult Organoid Growth Medium (osmolality, 270 mOsm/kg). 150 µL of Matrigel was added to the suspension. 50 µL of the mixture was mounted and spread on the glass region of a 35 mm glass bottom dish. The mixture was gelatinized by incubation for 10 min at 37°C. 1 mL of IntestiCult Organoid Growth Medium was added to the dish carefully. For clearing, the culture medium was replaced with SeeDB-Live/IntestiCult Organoid Growth Medium (refractive index 1.363) 2-3 hours before imaging. Phase contrast images were taken with an inverted microscope (DMI600B, Leica) equipped with a 10x NA0.4 dry objective lens. Fluorescence of enteroendocrine cells (EECs) in an organoid was imaged with an inverted confocal microscopy (TCS SP8, Leica) equipped with 20x NA0.75 multi-immersion lens and LASX software (Leica Microsystems).

Immersion was water and TDE/ddH_2_O (refractive index 1.366) for controls and SeeDB-Live samples, respectively. The correction collar was turned to the appropriate position. 488 nm laser was used to excite GCaMP6s expressed in EECs. To measure the Ca^2+^ responses of EECs, KCl (+30 mM at final concentrations) was added to the medium during imaging.

### Imaging of ES-cell-derived organoids

Mouse ES cells were maintained as described in the previous study ^27^. The cell line used in this study is a subline of the mouse ESC line, EB5 (129/Ola), in which the GFP gene was knocked in under the Rax promoter and Lifeact-mCherry gene was knocked in into Rosa26 locus. The cell line was provided by Dr. M. Eiraku at Kyoto University.

Cells were maintained as described in the previous study ^27^. For maintenance, cells were cultured in a gelatin-coated 100 mm dish. The dish contained maintenance medium, to which 20 µl of 106 units/ml LIF (Sigma-Aldrich) and 20 µl of 10 mg/ml Blasticidin (14499, Cayman) were added. Cells were incubated at 37 °C in 5% CO_2_. The maintenance medium consisted of Glasgow’s Modified Eagle Medium (G-MEM; 078-05525, Wako) supplemented with 10% Knockout Serum Replacement (KSR; 10828028-028, GIBCO), 1% Fetal Bovine Serum (FBS; GIBCO), 1% Non-essential Amino Acids (NEAA; 139-15651, Wako), 1 mM pyruvate (190-14881, Wako), and 0.1 mM 2-Mercaptoethanol (2-ME; M6250, Sigma-Aldrich). The solution was filtered through a 0.2 μm filter bottle, stored at 4 °C, used within one month.

For organoid induction, the serum-free floating culture of embryoid body-like aggregates with quick reaggregation (SFEBq) culture method was performed as described in study ^27^. In this method, 3,000 cells were suspended in 100 µl of differentiation medium in each well of a 96-well plate on day 0. On day 1, Matrigel (354230, Corning) was mixed with the differentiation medium and added to each well to reach a final concentration of 2.0%. This plate was incubated at 37 °C in a 5% CO_2_ environment. The differentiation medium consisted of G-MEM supplemented with 1.5% KSR, 1% NEAA, 1% pyruvate, and 0.1% 0.1 M 2-ME. This solution was filtered through a 0.2 μm filter bottle, stored at 4 °C, and used within one month.

For imaging, the Matrigel surrounding the organoids was reduced by pipetting gently in advance. Then the organoids were transferred from 96-well plate to 35 mm glass bottom dish (D11130H, Matsunami) coated with 0.1% (w/v) poly-L-lysine solution in H_2_O (P8920, Sigma-Aldrich) and 2.5 mg/ml Cell-Tak (354240, CORNING). The organoids were attached to the bottom by removing the medium as much as possible and incubating for 20 min at 37°C in a 5% CO_2_ environment. Images were captured using a LSM800 (ZEISS) equipped with a 25x NA0.8 multi-immersion lens. Immersion was water and TDE/ddH_2_O (refractive index 1.366) for controls and SeeDB-Live samples, respectively. On day 9 in SFEBq culture, 145 images were taken for each organoid at different *z* positions with 3 µm intervals within 432 µm. Organoids were incubated for 2 hours in SeeDB-Live medium adjusted at 270 mOsm/kg. Small incisions were made in the organoid by randomly inserting a glass capillary five times to facilitate penetration of SeeDB-Live into the internal vesicle.

Cultures for cortical organoid induction were performed as described in the study ^55^. In this method, the cortical organoid differentiation medium consisted of G-MEM supplemented with 10% KSR, 1% NEAA, 1% pyruvate, and 0.1% 0.1M 2-ME. The solution was filtered through a 0.2-μm filter bottle, stored at 4°C, and used within 1 month. On day 0, 3,000 cells were suspended in 100 µl of differentiation medium and placed in each well of a 96-well plate. The plates were incubated at 37°C in a 5% CO_2_ environment. On day 7, the aggregates were transferred to 35-mm bacterial-grade dish containing DMEM/F-12 with Glutamax (10565, Invitrogen) supplemented with N2 (17502-048, Invitrogen) and incubated in a 5% CO_2_, 40% O_2_ environment at 37°C. The medium was changed every 3 days. For Ca^2+^ imaging, the organoids were incubated with 5 µM Calbryte 630 (20721, AAT Bioquest) for 1 hour, transferred to 35 mm glass bottom dish and covered with cover glass (Matsunami). Images were captured using a LSM800 (ZEISS) equipped with a 25x NA0.8 multi-immersion lens. Immersion was water and TDE/ddH_2_O (refractive index 1.366) for controls and SeeDB-Live samples, respectively. On day 36 in SFEBq culture, 145 images were taken for each organoid at different *z* positions with 3 µm intervals within 432 µm. Organoids were incubated for 1 hour in SeeDB-Live medium. **Production of adeno-associated virus**

AAV-DJ-syn-jGCaMP8m-WPRE vector was generated using pGP-AAV-syn-jGCaMP8m-WPRE (Addgene #162375), pHelper (AAVpro Helper-free system, Takara), pAAV-DJ (Cell Biolabs), and the AAVpro 293T cell line (#632273, Takara) following the manufacturers’ instructions. Transfection was performed with PEIMAX (#24765-1, PSI). AAV vectors were purified using the AAVpro Purification Kit All Serotypes (#6666, Takara). AAV.PHP.S-CAG-jGCaMP8f-WPRE vector was generated using pAAV-CAG-jGCaMP8f-WPRE, pHelper, pUCmini-iCAP-PHP.S (Addgeen #103006), and the AAVpro 293T cell line as described previously ^56^. Briefly, the conditioned medium containing AAV vectors was filtered with a syringe filter to remove cell debris at 6 days post-transfection. The filtered medium was concentrated and formulated with D-PBS (-) using the Vivaspin 20 column pretreated with 1% BSA in PBS. Viral titers were measured using AAVpro Titration Kit (#6233, Takara) or THUNDERBIRD SYBR qPCR Mix (QPS-201, TOYOBO) with StepOnePlus system (ThermoFisher) or QuantStudio3 real-time PCR system (Applied Biosystems).

### Imaging of primary cardiomyocytes

Primary cultures of cardiomyocytes were prepared from P0 ICR mice as previously described^57^. The pups were anesthetized on ice and decapitated. The hearts were dissected and washed in PBS (-) containing 20 mM 2,3-butanedione monoxime (#B0753, Sigma). The hearts were cut into 0.5-1 mm pieces in Hanks’ Balanced Salt solution (HBSS (-); #084-08345, Fujifilm) containing 0.08% Trypsin-EDTA and 20 mM 2,3-butanedione monoxime and shaking at 4°C for 2 hours. L15 medium (#128-06075, Fujifilm) containing 1.5 mg/mL collagenase/dispase mix (#10269638001, Roche) and 20 mM 2,3-butanedione monoxime was added. 30 min after shaking at 37°C, the suspension was filtered through a 70 μm cell strainer. The trapped heart tissues were transferred to an L15 medium containing 1.5 mg/mL collagenase/dispase mix and 20 mM 2,3-butanedione monoxime and incubated for 10 min at 37°C. The suspension was filtered through the cell strainer again. After centrifugation at 100 g for 5 min, the pellet was resuspended with DMEM (high glucose) supplemented with 1% Penicillin-Streptomycin Solution and 10% FBS. The suspension was mounted on a cell culture dish for 2 hours. This helped the removal of highly adhesive cells. After gentle pipetting, the suspension was collected and plated on 35 mm dishes at 1.2 × 10^5^ cells/cm^2^. The cells were cultured in DMEM (high glucose), without phenol red and glutamine (#040-30095, Fujifilm) supplemented with 1% Penicillin-Streptomycin Solution, 10% FBS, and 1% glutamine at 37°C, 5% CO_2_. On day 1 *in vitro* (DIV-1), AAV.PHP.S-CAG-jGCaMP8f-WPRE was added at 2 × 10^10^ gc/mL. On DIV2, the culture medium was exchanged. The spontaneous activity of cardiomyocyte aggregates was measured with Leica TCS SP8 equipped with a 20x NA0.75 multi-immersion lens and LASX software (Leica Microsystems) at DIV-3 to 5. Phase contrast images were taken with an inverted microscope (DMI600B, Leica) equipped with a 10x NA0.4 objective lens.

### Imaging of primary hippocampal neuron culture

Primary cultures of hippocampal neurons were prepared from embryonic day 16 ICR mice. The embryos were taken from the uterus and decapitated in cold HBSS (-). The brain was extracted and put into a cold dissection medium consisting of HBSS (-) supplemented with 20 mM HEPES and 1% Penicillin/Streptomycin solution. The hippocampus was extracted from the brain and transferred to the dissection medium in a 15 mL tube. 2 mg/mL papain (#LS003119, Worthington) / HBSS (-) was activated for 5 min at 37°C. After filtration, the hippocampi were transferred to papain/HBSS (-) and incubated for 20 min at 37°C. 1 mL of 150 mg/mL Dnase I (#11284932001, Roche)/ HBSS (-) was added to the papain/HBSS (-) containing the hippocampi. The hippocampi were incubated for 5 min at 37°C. The hippocampi were washed twice with 2 mL of HBSS (-). The supernatant was replaced with 2 mL of Neurobasal medium (#21103-049, ThermoFisher) supplemented with 2% B27(#17504-044, ThermoFisher), 1% GlutaMax (#35050-061, ThermoFisher), and 1% Penicillin/Streptomycin solution. The cells were dissociated with gentle pipetting with a Pasteur pipette (Iwaki). The cells were then plated on a poly-D-lysine (#P7886, Sigma) coated 35-mm glass bottom dish at 1.5×10^5^ cells on a 12-mm-diameter coverslip and cultured in 5% CO_2_ at 37°C. On DIV-2, AAV-DJ-hsyn-jGCaMP8m-WPRE was added at 7 × 10^10^ gc/mL after half of the culture medium in the dishes was transferred to a 50 mL tube. Twenty-four hours after infection, the medium in the dishes was replaced with the culture medium kept in the 50mL tube together with the same amount of fresh medium. On DIV-7, half of the culture medium was transferred to a 50 mL tube. SeeDB-Live (refractive index 1.363) was made from this culture medium together with the same amount of fresh medium. The culture medium in the dishes was then replaced with SeeDB-Live. The spontaneous activity was measured with Leica TCS SP8 equipped with a 20x NA0.75 multi-immersion lens and LASX software (Leica Microsystems) on DIV-8 to 10. Phase contrast images were taken with an inverted microscope (DMI600B, Leica) equipped with a 20x NA0.7 objective lens.

### Confocal and two-photon imaging of acute brain slices

Thy1-GCaMP6f mice (P11-15) were used for Ca^2+^ imaging of the acute olfactory bulb slices. Thy1-YFP-H mice (P17-22) were used for morphological analyses. Mice were sacrificed with an overdose i.p. injection of pentobarbital (100-150 mg/kg) and decapitated. The brain was immediately harvested and placed in cold and O_2_-saturated artificial corticospinal fluid (ACSF: 125 mM NaCl, 3 mM KCl, 1.25 mM NaH_2_PO_4_, 2 mM CaCl2, 1 mM MgCl_2_, 25 mM NaHCO_3_, and 25 mM glucose). The brain was mounted on a silicone rubber block (Togawa rubber) and sliced using a microslicer (Dosaka EM) at 300 μm thickness. The slices were placed on a custom-made silicone chamber for imaging using an upright microscope as previously described ^31, 58^. The slices were recovered under the perfusion of O_2_-saturated ACSF for 1 hour at room temperature. For clearing, the brain slices were perfused with SeeDB-Live for 1 hour. To remove SeeDB-Live from tissue, ACSF was perfused for >1.5 hours.

The chamber was set under FV1000MPE microscope (Olympus/Evident) with Fluoview FV10-ASW software (Olympus/Evident, RRID:SCR_014215) and a 25x NA 1.05 water-immersion objective lens (Olympus/Evident, XLPLN25XWMP). Immersion was water and TDE/ddH_2_O (refractive index 1.366) for controls and SeeDB-Live samples, respectively. The correction collar was turned to the appropriate position. For one photon confocal imaging, a 473 nm laser was used. For two-photon imaging, a femtosecond laser (InSight DeepSee, SpectraPhysics) was tuned to 920 nm. For stimulation, 100 μM NMDA (Nacalai, Cat #22034-1) and 40 μM Glycine (Sigma, #G7126-100G) in ACSF or SeeDB-Live/ACSF was applied during Ca^2+^ imaging. Image data were analyzed with ImageJ. Briefly, small drifts were corrected by the Image Stabilizer plugin for Image J (https://imagej.net/plugins/image-stabilizer) when necessary. ROIs were created manually except for Fig. 2d and e, where ROIs were created by Cellpose. After fluorescence intensity was obtained, the data were analyzed with MATLAB software (MathWorks). The F_0_ was calculated by temporal median filtering (10 frames window). After the signal was filtered with temporal median filtering (3 frames window), the “findpeaks” function was applied for peak detection.

### Two-photon shadow imaging of acute brain slices

Acute brain slices were prepared from P18 wild-type C57BL/6N mice. The slice was mounted on a custom-made silicone chamber for imaging using an upright microscope as previously described ^31, 58^. The slice was perfused with ACSF at room temperature for 30 min for recovery. 40 μM Calcein was added to the ACSF. The slice was perfused with ACSF containing 40 μM Calcein for 1 hour. For clearing, the slice was perfused with SeeDB-Live containing 40 μM Calcein for 1 hour. For imaging, two-photon microscopy (MM201, Thorlabs) equipped with a 25x NA 1.05 water-immersion objective lens (Olympus/Evident, XLPLN25XWMP) and ThorImageLS software (Thorlabs) was used. A 920 nm femtosecond laser (ALCOR 920-4 Xsight, SPARK LASERS) was used. The images were averaged 5 times during the imaging.

### Electrophysiology

C57BL/6J mice were purchased from Japan SLC. Recordings were performed at P14 to 18. Mice were deeply anesthetized with isoflurane. Brains were quickly removed from mice and put into ice-cold ACSF bubbled with 95% O_2_ and 5% CO_2_. 300-µm thick acute coronal slices containing the primary somatosensory cortex were prepared using a vibratome (VT1200S, Leica). The slices were recovered in ACSF for at least 1 hour at room temperature (23-24°C) before recording for the control condition. For the SeeDB-Live condition, following the recovery in ACSF for 1 hour, the slices were transferred into SeeDB-Live solution saturated with 95% O_2_ and 5% CO_2_ for 1 hour at room temperature.

The slices were perfused with ACSF or SeeDB-Live at room temperature during the recording. Neurons were visualized by an infrared difference interference contrast (IR-DIC) video microscope with a 60x, 1.0 NA water immersion lens. Patch pipettes (4.6–9.9 MΩ) were filled with 130 mM potassium gluconate, 8 mM KCl, 1 mM MgCl_2_, 0.6 mM EGTA, 10 mM HEPES, 3 mM Na_2_ATP, 0.5 mM Na_2_GTP, 10 mM Tris_2_-phosphocreatine, and 0.2% biocytin (pH was adjusted to 7.35 with KOH and osmolality was adjusted to 295 mOsm/kg). Whole-cell recording was performed for layer 5b pyramidal neurons (Thick-tufted L5PT neurons) in the somatosensory area, which had large cell bodies and were located lower than 35 µm from the slice surface. Recordings were performed using MultiClamp700B amplifiers (Molecular Devices), filtered at 10 kHz using a Bessel filter and digitized at 20 kHz with Digidata 1440A digitizer (Molecular Devices), and stored using pClamp10 (Molecular Devices). Membrane potentials were given without correction for the liquid junction potential. A series resistance compensation was not used for recordings. When the series resistance exceeded 35 MΩ, the data were discarded. Data were analyzed using MATLAB.

To characterize firing properties, hyperpolarizing and depolarizing square current pulses were injected under current-clamp mode (+50 pA increment, 1 s, Fig. 3a). In characterizing membrane properties, the membrane potential was clamped at −60 mV, and square pulses (−5 mV, 50 ms) were applied in voltage-clamp mode (Supplementary Fig 6c). Spontaneous excitatory postsynaptic currents (sEPSCs) were recorded at the holding potential of −60 mV. Transient negative current responses with a peak amplitude of <−10 pA were detected as sEPSCs. The morphologies of the recorded neurons were visualized by staining biocytin with streptavidin-Cy3 (1:1000, #S6402; Sigma-Aldrich) after recording. Fluorescence images were obtained using confocal microscopy (LSM900, Zeiss) equipped with a 10x objective lens.

### *In utero* electroporation

*In utero* electroporation was performed as described previously ^31^. To label mitral cells at E12, 1 μg each of pCAG-GCaMP6f and pGP-pcDNA3.1 Puro-CAG-Voltron2 (Addgene #172909) plasmids were injected into the lateral ventricle. Electric pulses (a single 10-ms poration pulse at 72 V, followed by five 50-ms driving pulses at 40 V with 950-ms intervals) were delivered along the anterior-posterior axis of the brain with forceps-type electrodes (3-mm diameter, LF650P3, BEX) and a CUY21EX electroporator (BEX).

For imaging of the soma of L2/3 neurons in S1, 1 μg of pCAG-GCaMP6f and pGP-pcDNA3.1 Puro-CAG-Voltron2-ST (Addgene #172910) plasmids were injected into the lateral ventricle at E15. For imaging of the dendrite of L2/3 neurons in S1, 0.2 μg of pCAG-tTA2, 0.5 μg of pBS-TRE-mTQ2 and 1 μg of pGP-CAG-Voltron2 plasmids were injected. Electric pulses (a single 10-ms poration pulse at 72 V, followed by five 50-ms driving pulses at 42 V with 950-ms intervals) were delivered toward the medio-lateral axis of the brain with forceps-type electrodes (5-mm diameter, LF650P5, BEX) and an electroporator (CUY21EX, BEX).

### Voltage imaging of acute brain slices

Voltage imaging with Voltron2 was performed in acute brain slices of P4 – 11 ICR mice subjected to *in utero* electroporation at E12. Mice were anesthetized in ice and decapitated. The brain was immediately harvested and placed in cold and O_2_-saturated ACSF. The brain was mounted on a silicone rubber block (Togawa rubber) and sliced using a microslicer (Dosaka EM) at 300 μm thickness. The slices were incubated in O_2_-saturated ACSF containing 50 nM Janelia Fluor HaloTag Ligand 549 (#GA1110, Promega) at room temperature for 1 hour. After placing on a custom-made silicone chamber, the slices were then washed under the perfusion of O_2_-saturated ACSF for 1 hour (2 hour for control experiment) at room temperature. For clearing, the brain slices were perfused with SeeDB-Live for 1 hour.

The chamber was set under FV1000MPE microscope (Olympus/Evident) with Fluoview FV10-ASW software (Olympus/Evident, RRID:SCR_014215) and a 25x NA 1.05 water-immersion objective lens (Olympus/Evident, XLPLN25XWMP). Immersion was water and TDE/ddH_2_O (refractive index 1.366) for controls and SeeDB-Live samples, respectively. The correction collar was turned to the appropriate position. For widefield imaging, mercury arc lamp was used. GFP (U-MNIBA3: Ex 470-495 nm, Di 505 nm, Em 510-550 nm) and RFP filter set (U-MWIG3: Ex 530-550 nm, Di 570 nm, Em >575 nm) were used for GCaMP and Voltron2_549_, respectively. Epifluorescence was imaged with a high-speed CCD camera (#MiCAM02-HR or MC03-N256, BrainVision) at 0.3-7 ms/frame. For two-photon imaging of GCaMP6f, a femtosecond laser (InSight DeepSee, SpectraPhysics) was tuned to 920 nm. Image data were analyzed with ImageJ. Small drifts were corrected by the Image Stabilizer plugin. ROIs were created manually. After fluorescence intensity was obtained, the data were analyzed with MATLAB software (MathWorks). The F_0_ was calculated as temporal median filtering (100 frames window). For peak detection, the “findpeaks” function was applied. Threshold was set as Mean + 3×SD. Voltage changes within a mitral cell (single trial data) were visualized with BV Workbench (Brain Vision). 2D mean filter (3×3 pixels), drift removal (polyfit, degree 3) and Savitzky-golay filter (32 points, window size 5 frames) were applied. For the spike triggered averaged movie, 65 events of backpropagating action potentials were averaged based on the spike timing at the soma.

### Evaluation of BSA permeability in the brain *in vivo*

Adult mice (3 weeks of age) were used. Surgery and imaging were performed under ketamine (Daiichi-Sankyo) – xylazine (Bayer) (80 mg/kg and 16 mg/kg, respectively) anesthesia. During surgery and imaging, the depth of anesthesia was assessed by the toe-pinch reflexes, and supplemental doses were added when necessary. A head holder for mice (#SGM-4, Narishige) was used for surgery and imaging. Body temperature was maintained with a heating pad (Akizuki, M-08908). For the imaging, a craniotomy (∼5 mm in diameter) was made over S1 using a dental drill with a 0.5 mm drill tip. A silicone sealant, Kwik-Sil (#KWIK-SIL, WPI), was mounted surrounding the craniotomy. The dura matter was carefully removed by a micro hook (durotomy) (#10065-15, Muromachi Kikai). ACSF-HEPES (145 mM NaCl, 5 mM KCl, 2 mM CaCl_2_, 1 mM MgCl_2_, 10 mM HEPES, pH7.3) was used to prepare the control ACSF and SeeDB-Live/ACSF (refractive index 1.366). 1 mg BSA-CF594 (#20290, biotium) was dissolved in 100 μL SeeDB-Live. 50-100 μL of SeeDB-Live (1% BSA-CF594) was mounted onto the brain surface. Mice were sacrificed with an overdose of pentobarbital i.p. (100-150mg/kg) and decapitated. The brain was embedded in OTC compound (#4583, Sakura Finetek) and frozen with liquid nitrogen. The frozen sections were made with a cryostat (#CM3050S, Leica) and immediately imaged with an inverted microscope (#DMI600B, Leica) equipped with a 5x NA 0.1 dry objective lens.

### *In vivo* imaging of the mouse brain

Anesthetized adult Thy1-YFP-H mice (1-6 month-old) were used for imaging. After a craniotomy and durotomy (∼5 mm in diameter) over S1, a silicone sealant rim was created around the window to maintain fluid over the brain surface. 50-100 μL of ACSF-HEPES or SeeDB-Live/ACSF-HEPES was applied onto the brain surface. After removing a silicone sealant rim, a circular coverslip (>10 mm diameter) was fixed with silicone sealant. One hour after application of ACSF-HEPES or SeeDB-Live/ACSF-HEPES, the fluorescence of layer 5 pyramidal neurons was imaged using a two-photon microscopy (#MM201, Thorlabs) equipped with 25x NA 1.05 water-immersion objective lens (Olympus/Evident, XLPLN25XWMP) and ThorImageLS software (Thorlabs). Immersion was water and TDE/ddH_2_O (refractive index 1.366) for controls and SeeDB-Live samples, respectively. The correction collar was turned to the appropriate position. A 920 nm femtosecond laser (ALCOR 920-4 Xsight, SPARK LASERS) was used.

### *In vivo* calcium imaging from the mouse primary visual cortex

Anesthetized adult C57BL6/J mice (∼7 week-old) were used for *in vivo* calcium imaging of the primary visual cortex (V1). Under isoflurane anesthesia (2%) together with calprofen (10 mg/kg), atropine (0.3 mg/kg), dexamethasone (2 mg/kg), craniotomy and durotomy was performed over the V1 (2.2 mm square). A half of the exposed cortex was covered with a piece of square coverslip (2 × 1 mm), and the other half was left directly contacted to an extracellular solution for a subsequent pipette insertion and SeeDB-Live/ACSF-HEPES application. Anesthesia was continued at a lower concentration of isoflurane (0.125–0.5%), supplemented with chlorprothixene (1 mg/kg, Sigma) ^59^. Body temperature was maintained at 37 °C using a feedback-controlled heating pad. Then, the mice were placed under a two-photon microscope (Bermago II, Thorlabs).

For *in vivo* two-photon calcium imaging, we used jGCaMP8m or a synthetic calcium sensor Cal-520 AM. jGCaMP8m expression was achieved by AAV DJ-Syn-jGCaMP8m-WPRE (4.1×10^14 gc/mL, 150 nL at depth of 350 μm, 3 sites 200 μm apart), and jGCaMP8m was excited at 980 nm (MaiTai DeepSee eHP, SpectraPhysics). Cal-520 was introduced by bolus loading ^60^. Cal-520 AM (AAT Bioquest) was dissolved in 4 μL DMSO containing 20% pluronic-127, and was diluted with HEPES based ACSF (NaCl 145 mM, KCl 5mM, CaCl_2_ 2mM, MgCl_2_ 1 mM, HEPES 10 mM) to a final concentration of 570 μM. The Cal-520 AM solution also included 50 μM Alexa594 (Sigma) to visualize dye loading pipettes (at 1070 nm excitation by Fidelity-2, Coherent). The pipette was inserted and advanced obliquely under the two-photon microscope, to target the depth of ∼420 μm under the coverslip. The same pipettes were also used to record visual response of local field potential at multiple sites along the pipette track, to ensure that the retinotopic position of the dye loading site was in the monocular zone of V1 ^59^. Cal-520 AM solution was pressure injected (150 mbar, 2 min), and after an incubation time of 1 hour, two-photon imaging experiments were performed. Two-photon excitation of Cal-520 was at 920 nm (MaiTai DeepSee eHP, SpectraPhysics).

Visual stimulation was presented at an LCD monitor (iPad 3/4 Retina Display, Adafruit, refresh rate 60 Hz, gamma corrected) which was placed contralateral to the craniotomy side, covering an angle of 100° horizontal and 80° vertical in the visual hemifield. The monitor was moved and placed at a position that could evoke maximal full-field calcium response.

The stimulation was controlled by MATLAB programs originally developed by Cortex Lab at University College London ^59^. Full-screen sinusoidal drifting gratings were presented randomly at one of the eight directions (0-315°, 45° step) together with a blank condition.

After obtaining visual response to the drifting gratings in the control condition of ACSF-HEPES (more than 15 repeats), solution under the objective lens (∼1 mL) was replaced with SeeDB-Live/ACSF-HEPES. After an additional 1 hour incubation when we confirmed that baseline brightness was increased for jGCaMP8m and visual response to the same set of drifting gratings was obtained in the presence of SeeDB-Live/ACSF-HEPES.

Calcium traces for each cell were computed offline using a python version of Suite2p ^54^. Region of interest (ROI) were drawn over cells, and fluorescence signal for each ROI was extracted. Neuropil contamination was not corrected, considering correction factor could be different in the presence and absence of SeeDB-Live Baseline florescence (F0) was defined as 25% percentile of the fluorescence signals in the sliding window of 30-0 sec before each specific time point.

Visual response for each cell was further analyzed using MATLAB. First, Visual responsiveness was evaluated by Kruskal Wallis test using calcium response at 8-direction plus the blank conditions. Orientation selectivity was judged by Kruskal Wallis test using response at 8-direction conditions.

For orientation selective cells, preferred orientation, orientation selective index and tuning width were computed as follows. Orientation response was fitted with the sum of two gaussian curves. The preferred direction was defined as an angle that has the peak of the fitted gaussian curve. The orientation selective index (OSI) was defined as the depth of modulation from the preferred orientation to its orthogonal orientation ^61^.

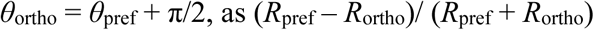

The tuning width was measured as full width at half maximum height of the fitted curve. The maximum response was average calcium signal at an orientation closest to the preferred orientation.

### Surgery for the mouse olfactory bulb

AAV1-Syn-FLEX-Volton2-ST-WPRE (Addgene # 172907-AAV1) was injected to 2-month-old Pcdh21-Cre mice for *in vivo* voltage imaging. Surgery was performed under ketamine (Daiichi-Sankyo) – xylazine (Bayer) (80 mg/kg and 16 mg/kg, respectively) anesthesia. A small circle (∼1 mm in diameter) was made with a dental drill over the right olfactory bulb. The AAV1 vector was injected into the center of dorsal olfactory bulb (200 µm depth), over a 12-min period, using a Nanoject III injector (Drummond Scientific Company). The total volume of the AAV solution was 120 nl.

Before imaging, mice were subjected to surgery under ketamine– xylazine (80 mg/kg and 16 mg/kg, respectively) anesthesia. After a craniotomy and durotomy over the right olfactory bulb, a silicone sealant rim was created around the window to maintain fluid over the brain surface. 50-100 μL of ACSF-HEPES or SeeDB-Live/ACSF-HEPES was applied onto the brain surface for one hour. After removing a silicone sealant rim, a circular coverslip (2 mm diameter) was mounted on the brain surface and fixed with a superglue and silicone sealant.

### *In vivo* calcium imaging in the mouse olfactory bulb

Anesthetized adult Thy1-GCaMP6f mice (2–4-month-old) were used for imaging. The fluorescence of mitral cells was imaged using FV1000MPE microscope (Olympus/Evident) with Fluoview FV10-ASW software (Olympus/Evident, RRID:SCR_014215) and a 25x NA 1.05 objective lens (Olympus/Evident, XLPLN25XWMP). Immersion was water or TDE/ddH_2_O (refractive index 1.366) for control or SeeDB-Live samples, respectively. The correction collar was turned to the appropriate position. Small drifts were corrected by the Image Stabilizer plugin. ROIs were created manually. After fluorescence intensity was obtained, the data were analyzed with MATLAB software (MathWorks). The ΔF was normalized to the mean intensity for 10 s before stimulus onset (F_0_), and the response amplitude was defined as the averaged ΔF/F_0_ during the first 10 s after stimulus onset.

### *In vivo* voltage imaging in the mouse olfactory bulb

Pcdh21-Cre mice injected with AAV1-Syn-FLEX-Volton2-ST-WPRE (Addgene #172907-AAV1) was anesthetized with ketamine and xylazine (80 mg/kg and 16 mg/kg, respectively). The mice were placed under FV1000MPE microscope (Olympus/Evident) and a 25x NA 1.05 water-immersion objective lens (Olympus/Evident, XLPLN25XWMP). Immersion was water and TDE/ddH_2_O (refractive index 1.366) for controls and SeeDB-Live samples, respectively. The correction collar was turned to the appropriate position. For excitation, mercury arc lamp was used. RFP filter set (U-MWIG3: Ex 530-550 nm, Di 570 nm, Em >575 nm) was used for Voltron2_549_-ST, respectively. Epifluorescence was imaged with a high-speed CCD camera (#MC03-N256, BrainVision) at 1-7 ms/frame.

Image data were analyzed using ImageJ. Small drifts were corrected using the Image Stabilizer plugin. ROIs were generated using a machine learning-based image segmentation tool (ilastik) ^62^. Further analysis was performed using MATLAB software (MathWorks).

ROIs smaller than 5 pixels and/or outside the glomeruli were excluded. F_0_ was calculated by temporal median filtering (25 frames window). The cross-correlation matrix was made from the time courses of each ROI. k-means clustering was applied with k = 11 (the number of glomeruli). To define subclusters, the spike synchronicity index between ROIs was calculated. Spikes were detected using the findpeaks function. Thresholds were set at Mean + 1.5×SD and Mean + 3×SD for the clustering and final spike detection, respectively. A synchronous event was defined when the spike in one ROI was detected within ± 7 ms (21 ms window) of the spike in another ROI. The synchronicity index was calculated by dividing the number of synchrony events by half of the total number of spikes in two ROIs. Hierarchical clustering was used to define subclusters. The threshold was set at 10% of the maximum distance in the dendrogram. To make a reference for the phase analysis, the averaged ΔF/F_0_ was calculated from all glomeruli. The ROIs for the glomeruli were created manually.

### Olfactometry

Odor stimulation using an olfactometer was described previously ^42^. The olfactometer consists of an air pump (AS ONE, #1-7482-11), activated charcoal filter (Advantec, TCC-A1-S0C0 and 1TS-B), and flowmeters (Kofloc, RK-1250). Valeraldehyde (TCI, Cat# V0001) and amyl acetate (FUJIFILM-Wako #018-03623) were diluted at 1% (v/v) in 1 mL mineral oil in a 50 mL centrifuge tube. Saturated odor vapor in the centrifuge tube was delivered to a mouse nose with a Teflon tube. The tip of the Teflon tube was located 2 cm from the nose of the animals. Diluted odors were delivered for 5 second at 1 L/min.

### *In vivo* awake imaging through a large cranial window with a plastic film

We made a large cranial window with a polyvinylidene chloride (PVDC) wrapping film, silicone plug, and a coverslip as described previously ^32^. Briefly, mice were injected with 15% mannitol solution (3 mL/100 g body weight; Sigma-Aldrich, M4125) and anesthetized with isoflurane (induction: 3%, surgery: 1%). We made 3 x 6 mm^2^ cranial window with scalpel (Kai, #11) over the right cortical hemisphere. Bleeding was stopped with a gelatin sponge (LTL pharma, Spongel). For clearing experiments, we applied SeeDB-Live/ACSF to the cranial window for 1 hour. SeeDB-Live/ACSF was replaced every 15 min. Then, a commercially available PVDC wrapping film (Asahi Kasei, Asahi Wrap or Saran Wrap, ∼11μm thick) with ∼1 mm margin was applied to the cranial window and firmly attached to the skull with superglue. A transparent silicone elastomer (GC, Exaclear) and a glass coverslip (Matsunami 18 x 18 No.1) were then applied on top of the wrap. The coverslip was then sealed with waterproof film. RCaMP3 was introduced to S1 by AAVs (AAV2/1-CAG-DIO-RCaMP3-WPRE + AAV1-hSyn-Cre.WPRE.hGH (Addgene, #105553-AAV1)) and imaged with two-photon microscopy (Movable Objective Microscope, Sutter) at 1040 nm excitation with InSight X3 (Spectra-Physics) and a 25x objective lens (Olympus, XLPLN25XWMP2). The cranial window with a plastic wrap was carefully removed with scalpel for repeated clearing experiments. Whisker stimulation was performed with air puffs produced by Picoxpritzer (Parker).

### *In vivo* voltage imaging from the mouse primary somatosensory cortex and olfactory bulb

For voltage imaging of L2/3 neuron somata from P17-30 ICR mice subjected to in utero electroporation at E15, surgery and imaging were performed under urethane (Sigma, #U2500) anesthesia (1.9 g/kg). For imaging of the dendrites from the 2-month-old ICR mice, surgery was performed under ketamine (Daiichi-Sankyo) – xylazine (Bayer) (80 mg/kg and 16 mg/kg, respectively) anesthesia. After a craniotomy and durotomy (3-5 mm in diameter) over S1, a silicone sealant rim was created around the window to maintain fluid over the brain surface. 50-100 μL of ACSF-Hepes containing 50 nM Janelia Fluor HaloTag Ligand 549 (#GA1110, Promega) was applied onto the brain surface for 1 hour. After removing the mounted solution, 50-100 μL of ACSF-HEPES was mounted for 1 hour (2 hour for control experiment). For clearing, SeeDB-Live/ACSF-HEPES was applied onto the brain surface for 1 hour. After removing a silicone sealant rim, a circular coverslip (>10 mm diameter) was fixed with superglue.

The mice were placed under FV1000MPE microscope (Olympus/Evident) and a 25x NA 1.05 water-immersion objective lens (Olympus/Evident, XLPLN25XWMP). Immersion was water and TDE/ddH_2_O (refractive index 1.366) for controls and SeeDB-Live samples, respectively. The correction collar was turned to the appropriate position. For excitation, mercury arc lamp was used. RFP filter set (U-MWIG3: Ex 530-550 nm, Di 570 nm, Em >575 nm) was used for Voltron2_549_, respectively. Epifluorescence was imaged with a high-speed CCD camera (#MC03-N256, BrainVision) at 1-3 ms/frame. Image data were analyzed with ImageJ. Small drifts were corrected by the Image Stabilizer plugin. ROIs were created manually. After fluorescence intensity was obtained, the data were analyzed with MATLAB software (MathWorks). The F_0_ was calculated as temporal median filtering (100 frames window). For movie, 3D median filter (1 pixel radius) was applied.

### Statistical analysis

MATLAB was used for statistical analysis. Sample sizes were not pre-determined. Number of animals was indicated within figure legends. All statistical tests were performed using two-sided tests. Student’s t-test was used in Fig. 3i, Supplementary Fig. 4d. Paired Student’s t-test was used in Figure 4f. Wilcoxon rank sum test was used in Fig. 2e, 3b, Supplementary Fig. 5g, 6d, e, g, i, 7d and e. Wilcoxon signed-rank test was used in Fig. 4k. Wilcoxon rank sum test corrected with Holm-Bonferroni correction was used in Fig. 1m, p, 3c, Supplementary Fig. 1f, 3f and g. Multiple comparison test with Bonferroni correction was used in Fig. 3e, f and Supplementary Fig. 7a and b. Dunnet’’s multiple comparison test was used in Fig. 1f, j, k, Supplementary Fig. 1a, g, 2a and b. In box plots, the middle bands indicate the median; boxes indicate the first and third quartiles; and the whiskers indicate the minimum and maximum values. Data inclusion/exclusion criteria are described in figure legends.

## Materials availability

New plasmids (pCAG-GCaMP6f and pAAV-CAG-jGCaMP8f-WPRE) generated in this study will be deposited to Addgene (https://www.addgene.org/Takeshi_Imai/).

## Data and code availability

Raw image data used in this study will be deposited to SSBD:repository (http://ssbd.qbic.riken.jp/set/2024xxxx/). Detailed protocols and technical tips will be described in SeeDB Resources (https://sites.google.com/site/seedbresources/). Requests for additional program codes and data generated and/or analyzed during the current study should be directed to and will be fulfilled upon reasonable request by the Lead Contact.

## Acknowledgements

We thank H. Zeng (*Ai162*), J. Sanes (Thy1-YFP-H), K. Svoboda (Thy1-GCaMP6f), and E. Deneris (*ePet-Cre*) for mouse strains; A. Miyawaki for cell lines (HeLa/Fucci2); M. Eiraku for ES cell lines (EB5); D. Kim & GENIE Project for pGP-CMV-GCaMP6f (Addgene plasmid # 40755); GENIE Project for pGP-AAV-syn-jGCaMP8m-WPRE, pGP-AAV-syn-jGCaMP8f-WPRE, pGP-pcDNA3.1 Puro-CAG-Voltron2 and pGP-pcDNA3.1 Puro-CAG-Voltron2-ST, pGP-AAV-syn-FLEX-Volton2-ST-WPRE (Addgene plasmid # 162375, 162376, 172909, 172910, 172907 and viral prep #172907-AAV1); V. Gradinaru for pUCmini-iCAP-PHP.S (Addgene plasmid # 103006); J. Wilson for AAV1-hSyn-Cre.WPRE.hGH (Addgene viral prep # 105553-AAV1); M-T. Ke and M. Morimoto for evaluating our earlier versions of the clearing medium; S. Uchida, K. Miyamichi, K. Yashiro, and S-H. Chou for sharing reagents; M. Nishihara and E. Nozoe for technical assistance. We also thank The Research Support Center, Research Center for Human Disease Modeling, Kyushu University Graduate School of Medical Sciences, which was in part supported by Mitsuaki Shiraishi Fund for Basic Medical Research. This work was supported by grants from CREST program (JPMJCR2021 to T.I.) of the Japan Science and Technology Agency (JST) (T.I.), AMED (JP23wm0525012 to TI, JP19dm0207080 to K.K., JP19dm0207079 to S.M), the JSPS KAKENHI (JP21H00205, and JP21H05696 to T.I., JP21H02140 and JP22K18373 to S.I., JP19K06886 to S.F., JP22K06446 and JP22H05094 to N.N-T., JP22H05161, JP22H00460, JP23K18161 to K.K., JP22H02718 to S.M.), World Premier International Research Center Initiative (WPI-PRIMe) (K.H.), the Mochida Memorial Foundation for Medical and Pharmaceutical Research, the Uehara Memorial Foundation (T.I.).

## Author Contributions

T.I. conceived the project. S.I. and T.I. designed the experiments. S.I. performed all the experiments for optimizing SeeDB-Live. N.N-T. and Y.T. performed electrophysiology experiments. Y.K., H.T., H.T. and T.K.S. performed *in vivo* imaging. S.M., K.K., and T.N. performed awake *in vivo* imaging with S.I. S.F. performed *in utero* electroporation. R.Y., A.T., M.M., Y.N., K.H., and S.O. performed organoid experiments. T.Y. and M.S. developed RCaMP3. T.I. supervised the project. S.I. and T.I. wrote the manuscript with inputs and feedback from all the authors.

## Supplementary Figures

**Supplementary Figure 1.**
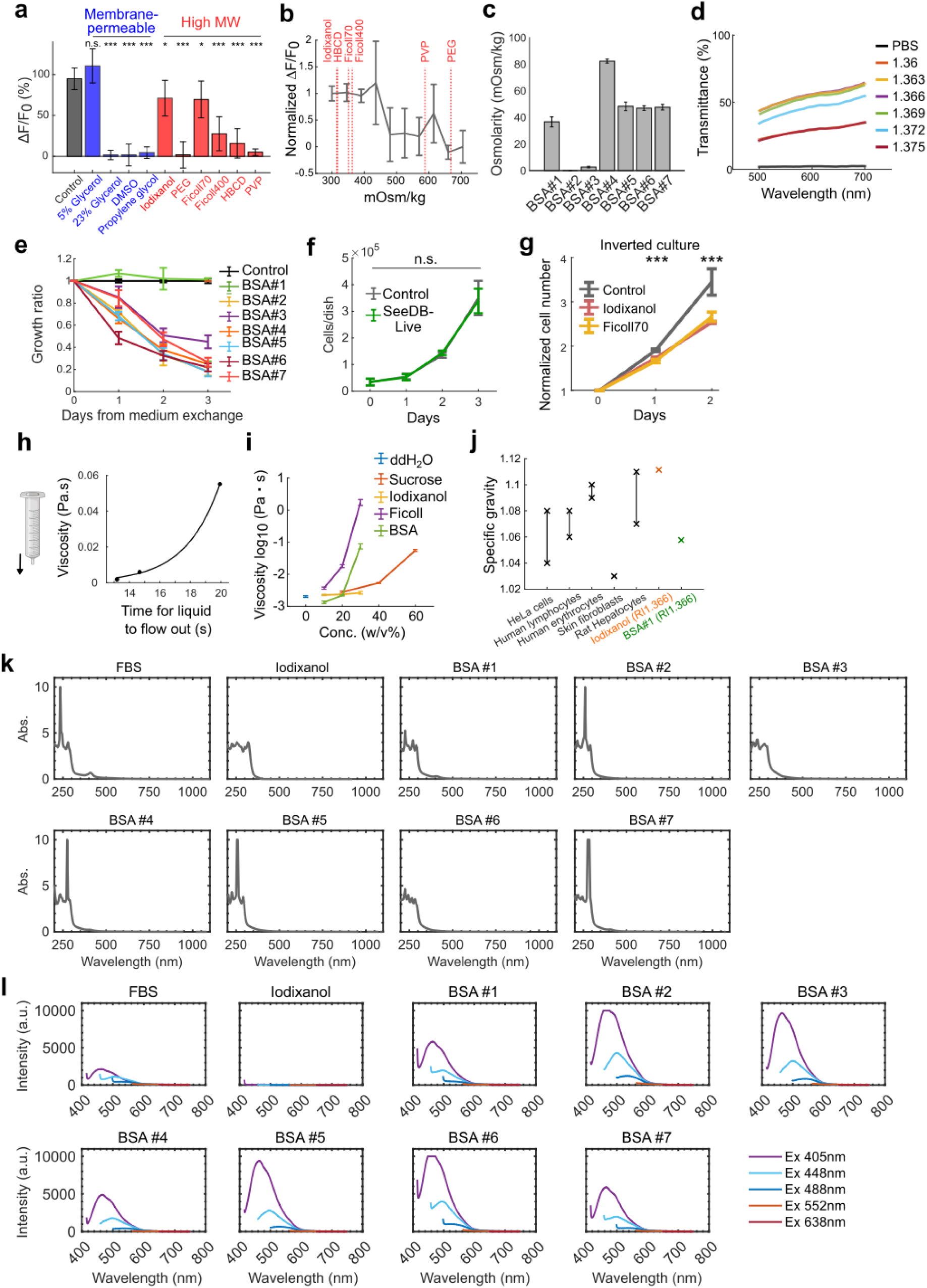
Evaluation of candidate chemicals. **(a)** GCaMP6f responses to 50 μM ATP measured with a plate reader. The refractive index was adjusted to 1.365 except for 5% glycerol (refractive index 1.348). Osmolarity was not adjusted to isotonicity (hypertonic, 300-700 mOsm/kg see also B) in this experiment. After incubation with the clearing media for 4 hours, GCaMP6f responses were measured. n = 8 wells. \*\*\**p* < 0.001; \*\**p* < 0.01; \**p* < 0.05; n.s., not significant (*p* ≥ 0.05) (Dunnet’’s multiple comparison test). **(b)** GCaMP6f responses to 50 μM ATP in the media with variable osmolality (300-700 mOsm /kg). The osmolality was adjusted with 10x PBS. The osmolalities of high MW media are also indicated. n = 8 wells. **(c)** The osmolality of different BSA products in aqueous solution (refractive index 1.365, in ddH_2_O). n = 3 each. There are several products of BSA with different grades of purity are available from different companies. The osmolality was slightly different among these products, suggesting that salts remain in some of the products. Osmolarities of low-salt BSA (#3) was 2.7 mOsm/kg, consistent with its molar concentration (2.3 mM). **(d)** Transmittance of HeLa cell suspension (4 × 10^6^ cells/mL) in isotonic BSA#1/PBS at refractive index 1.365-1.375. **(e)** Growth ratio in refractive index-optimized (refractive index 1.363, 320 mOsm/kg) medium of different BSA products. Cell numbers were counted based on the cell nuclei in the fluorescence images, and the number was normalized by the control data at each stage (n = 5 wells). Cell growth was lower for some BSA products. Variability among different BSA products may be due to the contamination of toxic chemicals. **(f)** Growth curve of HeLa cells cultured in a 35 mm dish. Cell number was measured with a hemocytometer after trypsinization. The medium was isotonic, and their refractive indices were adjusted to 1.363. n.s.; not significant (*p* ≥ 0.05) (Wilcoxon rank sum test corrected with Holm-Bonferroni correction). **(g)** Growth curve of HeLa/Fucci2 cells cultured in the inverted culture in 384-well plates. Cell number was counted based on fluorescence images of the nuclei. n = 3 dishes. \*\*\**p* < 0.001 (Dun’ett’s multiple comparison test). **(h)** Standard curve for the viscosity measurement using sucrose solution. Flow speed was determined using 50 mL syringes. Plot was fitted with a single-exponential curve. The viscosity of sucrose solution is after the previous literature ^52^. **(i)** Viscosity of candidate solutions (in ddH_2_O) based on the flow rate in **(h)**. n = 3 each. **(j)** Specific gravity of candidate solutions in PBS. Specific gravity of cells are cited from previous studies ^64^. **(k, l)** Some of the BSA products have autofluorescence signals of unknown origin, especially for UV to blue range; therefore, we carefully selected low autofluorescence and low toxicity products from multiple suppliers. Absorbance **(k)** and fluorescence **(l)** of fetal bovine albumin (FBS), iodixanol, and BSA from different manufacturers are shown. Except for FBS, they are adjusted to 15% (w/v) in ddH_2_O. FBS was non-diluted. BSA#1: BSA crystal (bioWORLD), BSA#2: BSA crystal (Fujifilm), BSA#3: BSA Low Salt (Fujifilm), BSA#4: BSA pH 5.2 (Fujifilm), BSA#5: BSA pH 7.0 (Fujifilm), BSA#6: BSA Globulin Free (Fujifilm), BSA#7: BSA Protease Free (Fujifilm).

**Supplementary Figure 2.**
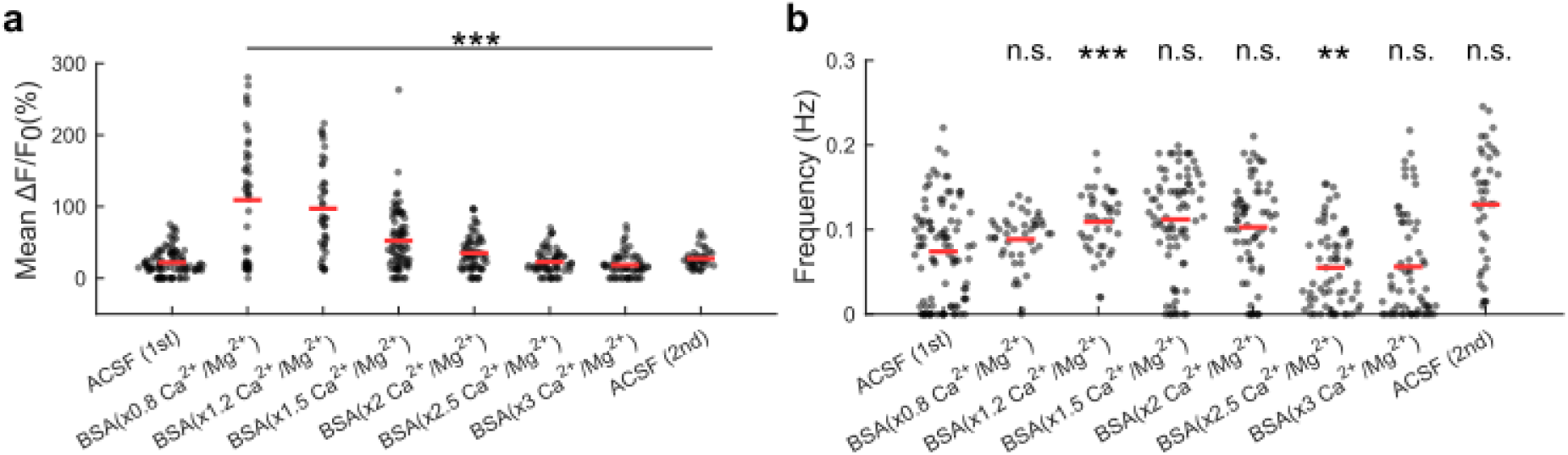
Optimization of divalent cations in SeeDB-Live. **(a, b)** Spontaneous activity of mitral cells in the olfactory bulb was measured under different concentrations of divalent cations. Amplitude **(a)** and frequency **(b)** were analyzed. Thy1-GCaMP6f mice (P11-14) were used for two-photon imaging of acute olfactory bulb slices. The ratio of added Ca^2+^ and Mg^2+^ was 2:1, following the formula of the ACSF (2 mM Ca^2+^ and 1 mM Mg^2+^). Normal calcium activity was observed at 2x higher concentrations of divalent cations (4 Ca^2+^ and 2 Mg^2+^). n = 95, 43, 40, 76, 65, 64, 63 and 40 cells from 3 slices. \*\*\**p* < 0.001; \*\**p* < 0.01; n.s., non-significant (*p* ≥ 0.05) (Dun’ett’s multiple comparison test).

**Supplementary Figure 3.**
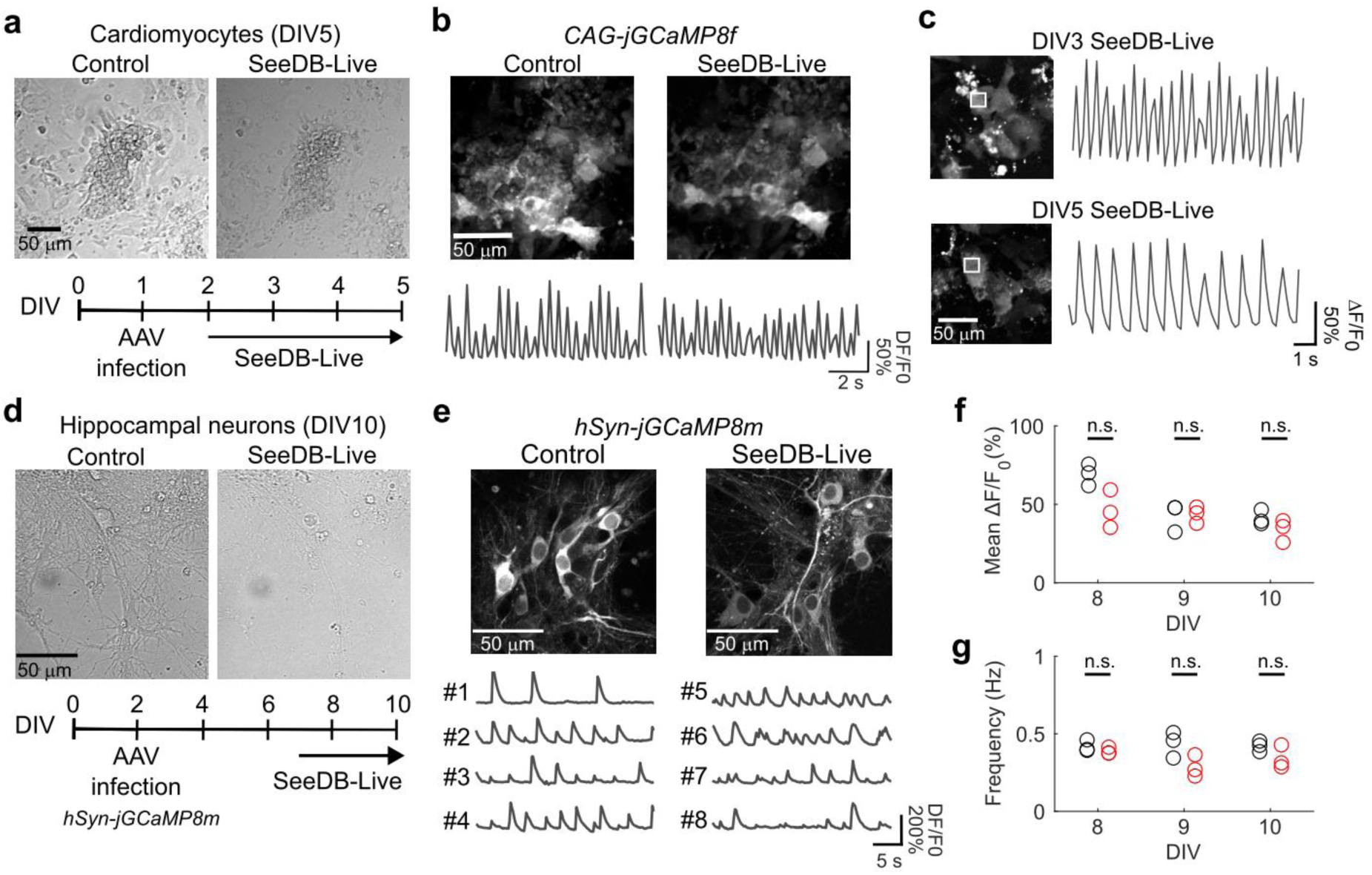
Long-term culture in SeeDB-Live. **(a)** Primary culture of cardiomyocytes. Phase contrast image (top) and the timeline of the culture (bottom). AAV.PHP.S-CAG-jGCaMP8f-WPRE (2 × 10^10^ gc/mL) was added at DIV1 and the medium was replaced at DIV2. Cells were cultured in SeeDB-Live from DIV2 to DIV5. **(b)** Primary culture of cardiomyocytes before and after the medium change to SeeDB-Live. Confocal image (top) and spontaneous calcium signals (bottom) at DIV2. **(c)** GCaMP8f signals of cultured cardiomyocytes at DIV3 and DIV5, showing spontaneous calcium signals. **(d)** Primary culture of hippocampal neurons. Phase contrast image (left) and the timeline of the culture (right). AAV-DJ-hSyn-jGCaMP8m-WPRE (7 × 10^10^ gc/mL) was infected at DIV2. Neurons were cultured in SeeDB-Live medium from DIV7. **(e)** GCaMP8m signals of cultured hippocampal neurons before and after the medium change to SeeDB-Live. **(f, g)** Amplitude and frequency of spontaneous calcium signals for the control (black) and SeeDB-Live (red). n = 3 dishes each. n.s., non-significant (Wilcoxon rank sum test).

**Supplementary Figure 4.**
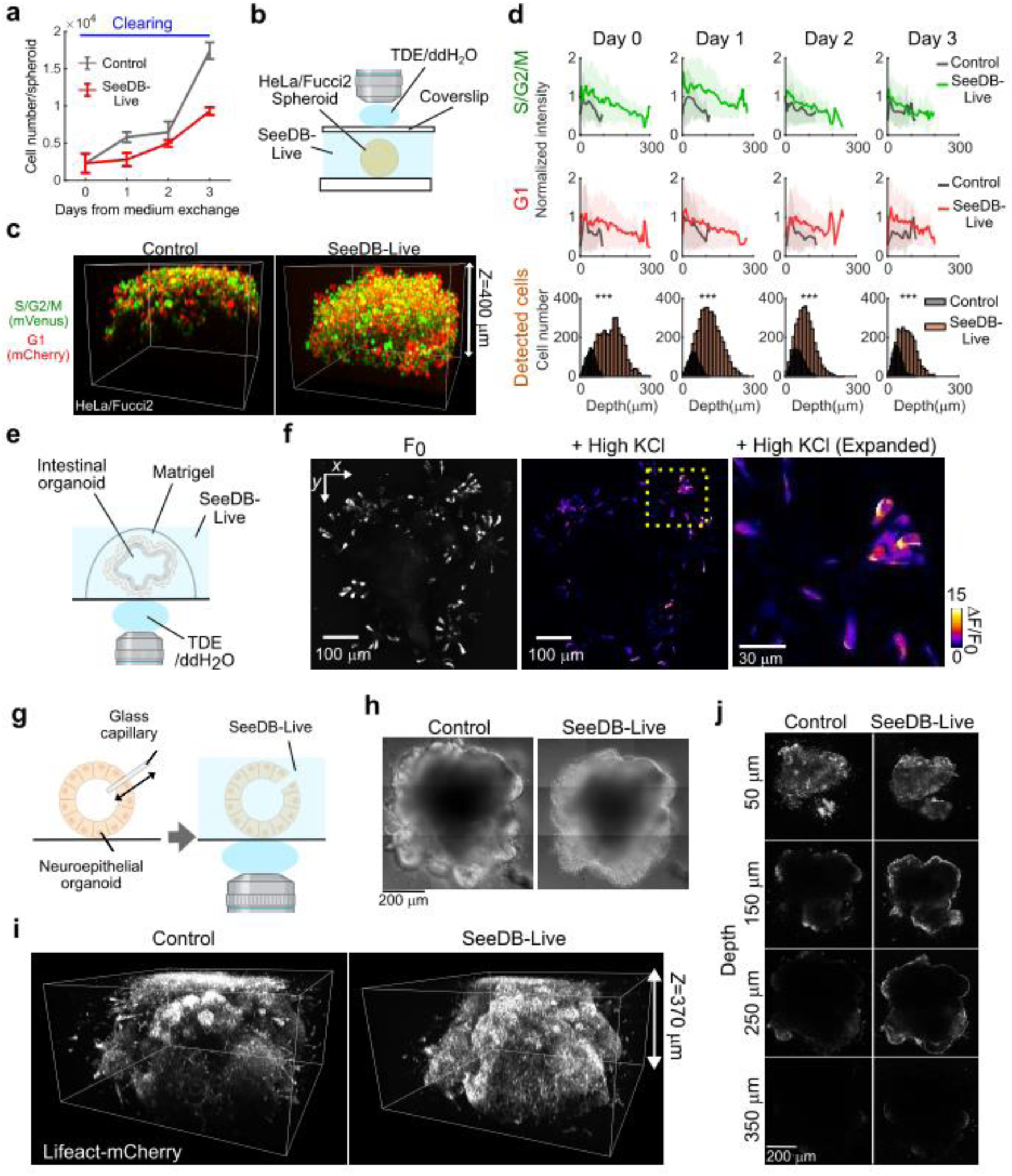
Clearing live spheroids and organoids with SeeDB-Live. **(a)** Growth of HeLa/Fucci2 spheroids cultured continuously in SeeDB-Live. Cell number was calculated with a hemocytometer after trypsinization. n = 3 spheroids each. Half of the medium was replaced daily. **(b)** Schematic diagram of fluorescence imaging of spheroids. TDE/ddH2O (refractive index 1.366) was used as immersion. The correction collar was optimized. **(c)** Three-dimensional fluorescence images of a HeLa/Fucci2 cell spheroid in the control and SeeDB-Live media (4 hour clearing per day as shown in Fig. 1m). See also Supplementary Video 1. **(d)** Depth-dependent fluorescence intensity from cell nuclei in the central portion of the spheroids (top, mVenus; middle, mCherry) and the number of cells detected in Cellpose (bottom). Fluorescence intensity indicate the mean intensity of all the cell nuclei in each *z* plane. \*\*\**p* < 0.0001 (Student’s t-test). **(e)** Schematic diagram of fluorescence imaging of intestinal organoids in Matrigel. TDE/ddH_2_O (refractive index 1.366) was used as immersion. The correction collar was in the optimal position. **(f)** Responses of enteroendocrine cells to high potassium stimulation (30 mM at final concentrations). GCaMP6s signals are shown for the intestinal organoids derived from ePet-Cre; Ai162 mice (EEC-GCaMP6s). F_0_ (left) and ΔF/F_0_ (right) images are shown. Magnified image of the inset is shown on the right. **(g)** ES cell-derived neuroepithelial organoids (day 9). The bright field images before and after SeeDB-Live treatment. **(h)** Fluorescence images of the Lifeact-mCherry-expressing neuroepithelial organoid at different depths before and after SeeDB-Live treatment. **(i)** 3D fluorescence images of Lifeact-mCherry-expressing neuroepithelial organoid before and after SeeDB-Live treatment. A representative sample out of three with similar results. Normal medium (left) and SeeDB-Live (refractive index 1.366; right). Small incision was made in the organoid before SeeDB-Live treatment.

**Supplementary Figure 5.**
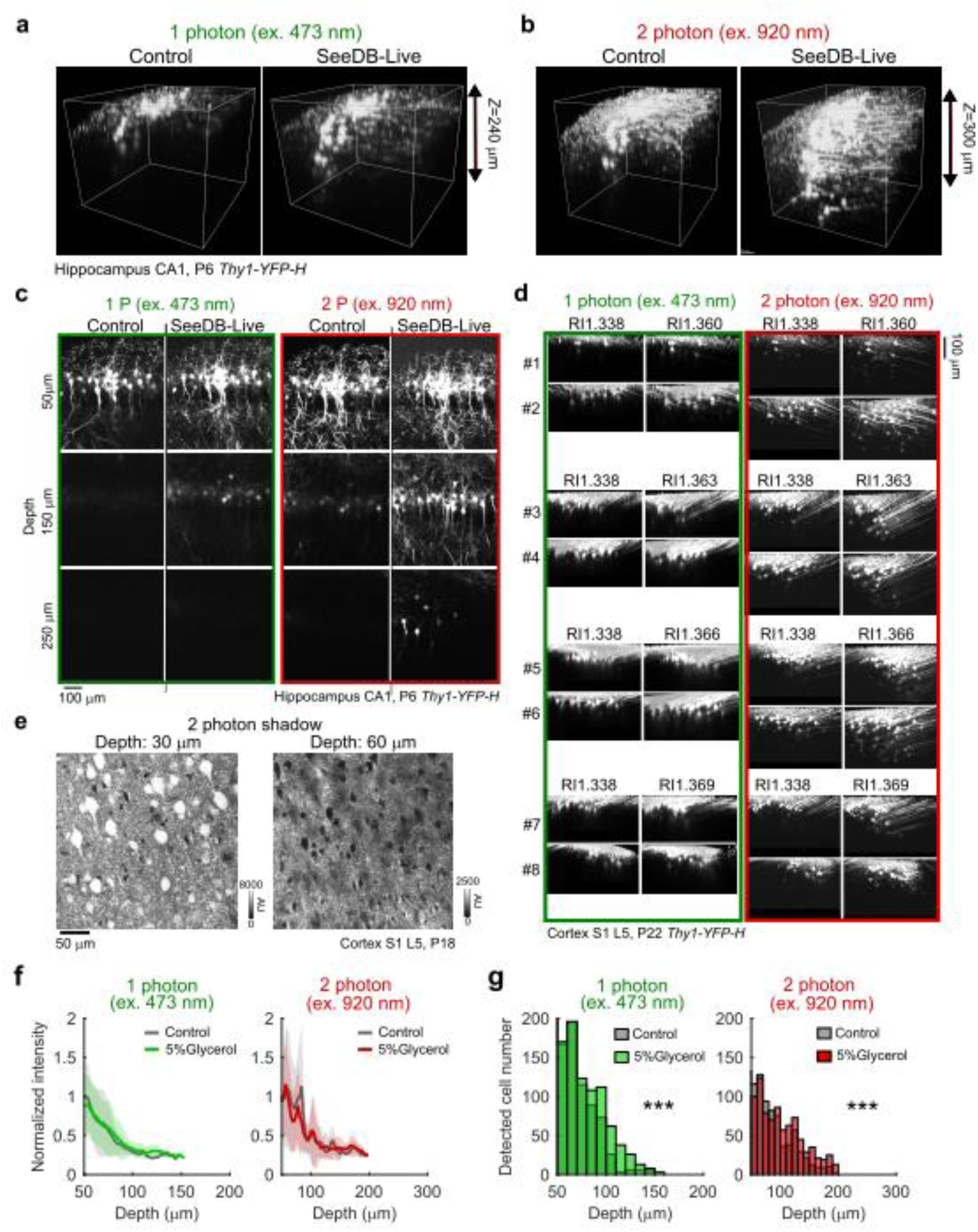
Clearing acute brain slices with SeeDB-Live. **(a, b)** Confocal **(a)** and two-photon **(b)** images of acute hippocampal slices cleared with SeeDB-Live (refractive index 1.363, 310 mOsm/kg in ACSF). Thy1-YFP-H mice (age P6) were used. **(c)** Fluorescence images of the hippocampus at different depths. **(d)** Fluorescence images at different refractive indices. Thy1-YFP-H mice (age, P22) were used. SeeDB-Live at different refractive indices (refractive index 1.338-1.369, 310 mOsm/kg) were tested. The optimal refractive index was 1.366, consistent with results from HeLa cells (Figure 1h). **(e)** Two-photon shadow images at the superficial region of an acute brain slice. The slice (S1 L5) was perfused with ACSF containing 40 μM Calcein. Bright signals are somata labeled with Calcein. The fluorescent dye was incorporated into the damaged cells at a depth of ∼30 μm. Bright dots (neurites of dead neurons) were still visible at a depth of ∼60 μm. **(f)** Fluorescence intensity from cell bodies in *x-y* fluorescence images of S1 L5 pyramidal neurons in 5% glycerol/ACSF. Acute brain slices of Thy1-EYFP-H mice (P19-21) were imaged before and after 5% glycerol/ACSF treatment. ROIs were generated by Cellpose, and mean fluorescence in ROIs are shown. **(g)** Number of detected cells in each depth in S1. Cells were detected by Cellpose. \*\*\**p* < 0.0001 (Wilcoxon rank sum test).

**Supplementary Figure 6.**
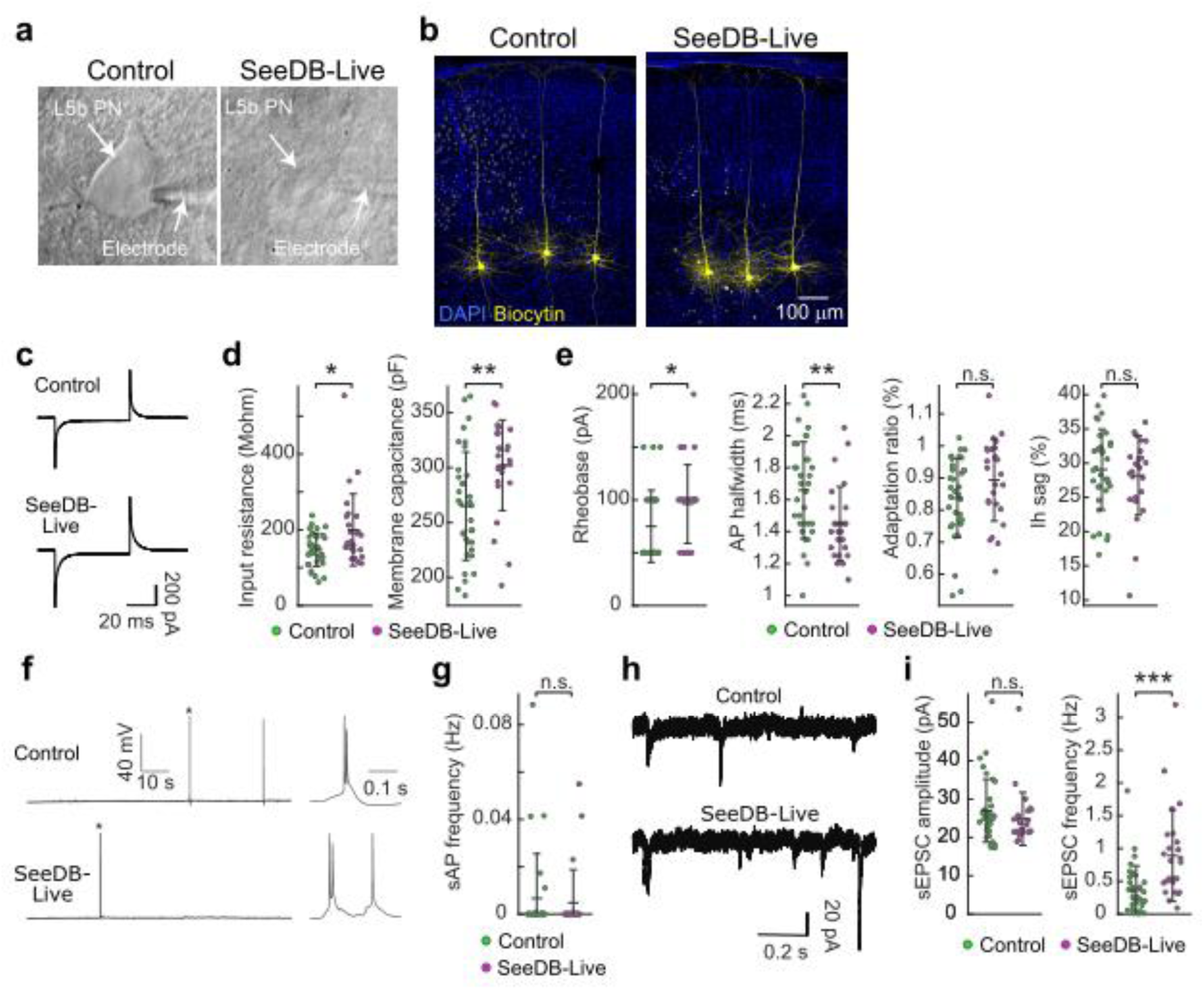
Electrophysiological properties of neurons under SeeDB-Live. **(a)** Infra-red differential interference contrast (IR-DIC) images of representative L5 pyramidal neurons under electrophysiological recording. As the tissue became transparent under SeeDB-Live, neurons were more difficult to identify using IR-DIC images. **(b)** Recorded neurons visualized by biocytin staining. They were all thick-tufted L5PT neurons. **(c)** Current responses to the test pulse (−5 mV, 50 ms). **(d)** Membrane properties, calculated from the current response to the test pulse. Individual data and mean ± SD are shown. **(e)** The minimum current needed to elicit an AP (Rheobase), firing properties (AP halfwidth and Adaptation ratio), and hyperpolarization-activated current (Ih)-related sag in the voltage response to a hyperpolarizing current. **(f)** Representative traces of spontaneous membrane potential fluctuation. Right; spontaneous AP (sAP) seen at the time shown on the left (*). **(g)** sAP frequency of individual neurons. **(h)** Representative traces of spontaneous current responses at the holding potential of −60 mV. **(i)** Spontaneous EPSC (sEPSC) amplitude and frequency. n = 34 for ACSF and 26 for SeeDB-Live. Mean ± SD are shown. \*\*\**p* < 0.001; \*\**p* < 0.01; \**p* < 0.05; n.s., not significant (*p* ≥ 0.05) (Wilcoxon rank sum test).

**Supplementary Figure 7.**
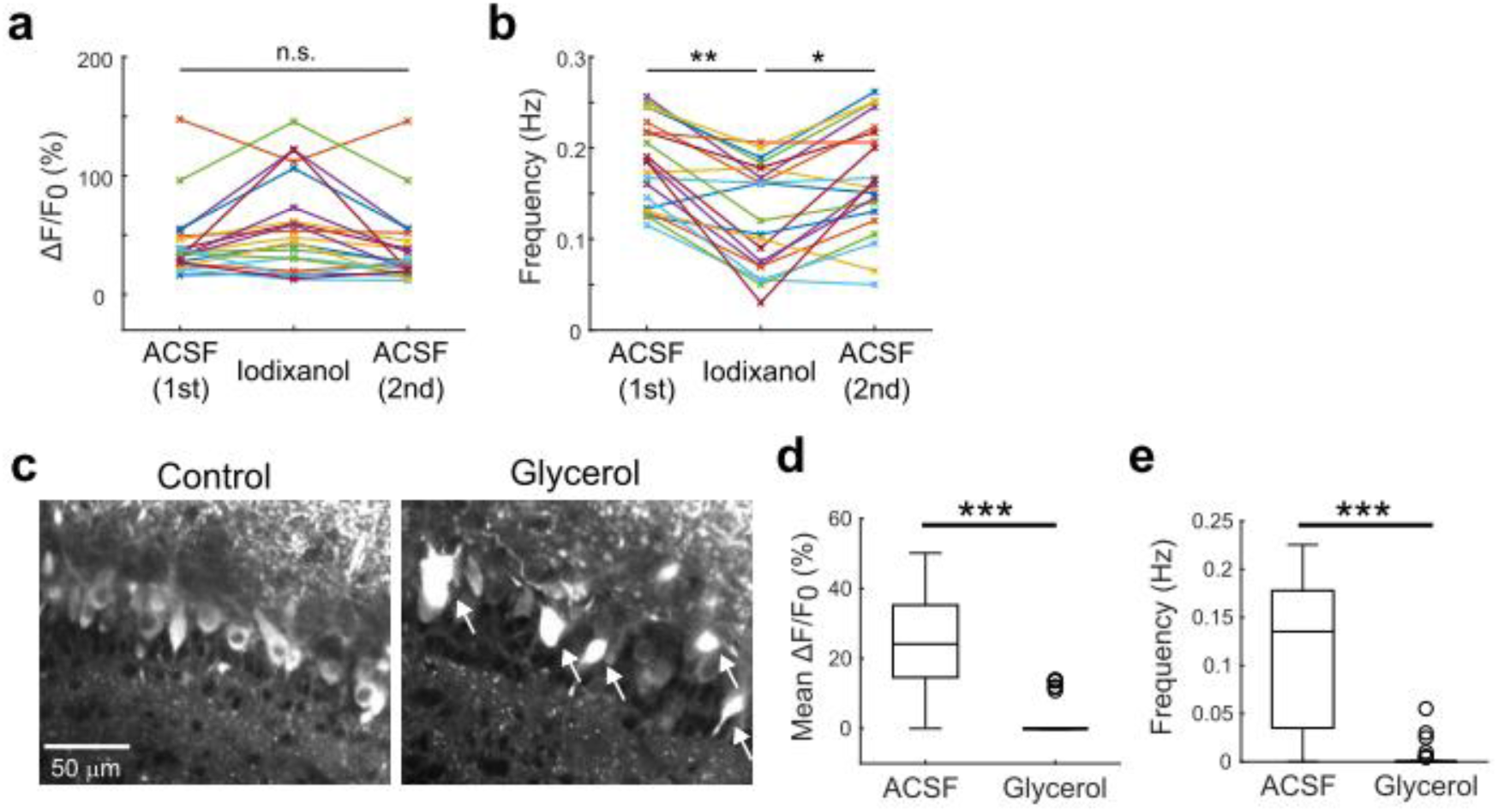
Evaluation of iodixanol and glycerol media for functional imaging in acute brain slices. **(a, b)** Amplitude and frequency of spontaneous activity in the same set of mitral cells in ACSF and Iodixanol/ACSF (RI1.366). n = 21 cells from 3 mice. *p < 0.05; **p < 0.01; n.s., non-significant (multiple comparisons with Bonferroni correction). **(c-e)** Spontaneous activity in the acute olfactory bulb slices was evaluated for glycerol-containing ACSF at a refractive index of 1.366. Thy1-GCaMP6f mice (age, P11) were used. Spontaneous activity was eliminated, and bright GCaMP6f signals started to appear 30 min after treatment with glycerol containing ACSF, suggesting that neurons are sick/dead in this condition **(c)**. Mean amplitude **(d)** and frequency **(e)** of spontaneous activity in mitral cells. \*\*\**p* < 0.001 (Wilcoxon rank sum test).

**Supplementary Figure 8.**
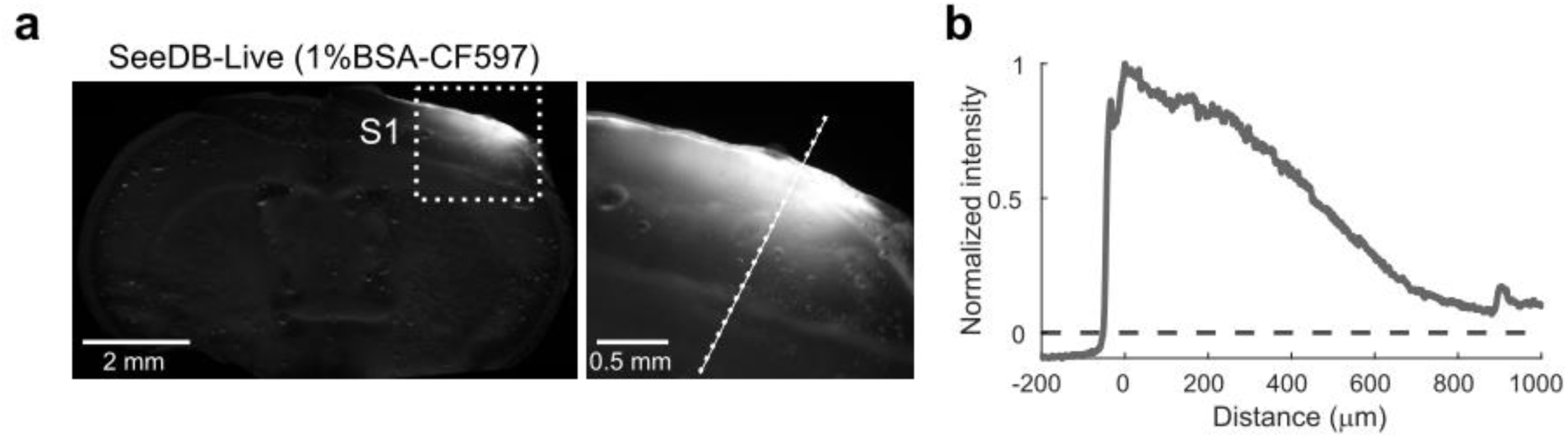
Penetration of BSA into the brain in live animals. **(a)** Penetration of BSA into the brain in live animals. After the craniotomy and durotomy, SeeDB-Live with 1% BSA-CF594 (refractive index 1.366, 310 mOsm/kg) was applied on the surface of S1. One hour later, the brain was isolated without perfusion and immediately frozen. Frozen sections were cut and imaged. **(b)** Line plot of BSA-CF594 fluorescence along the dotted line (from cortical surface) in **(a)**. BSA-CF594 penetrated up to a depth of ∼800 μm.

**Supplementary Figure 9.**
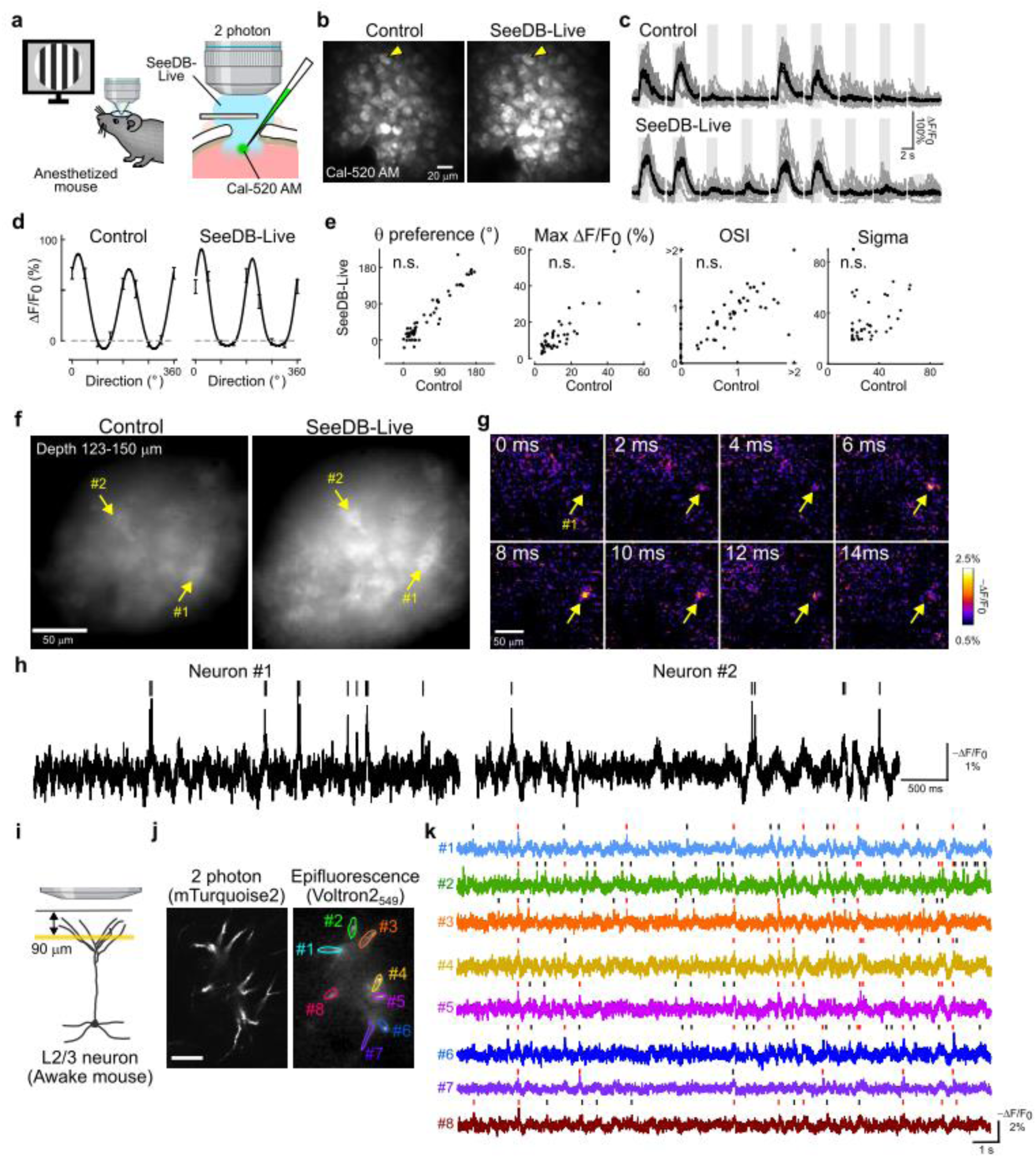
*In vivo* imaging in anesthetized and awake mice. **(a)** Calcium imaging of L2/3 neurons in the primary visual cortex (V1) before and after clearing with SeeDB-Live. In anesthetized animals, neurons were labeled with Cal-520-AM and imaged with two-photon microscopy. **(b)** Basal fluorescence of Cal-520 without visual stimulation. L2/3 neurons at a depth of ∼420 μm. **(c, d)** Responses of a representative L2/3 neuron (indicated by arrowheads in **h**) to visual grating stimuli before and after clearing with SeeDB-Live. **(e)** Preferred orientation, maximum responses (ΔF/F_0_), orientation selective index (OSI), and tuning width (Sigma) for the same set of L2/3 neurons (65 neurons in total) before (*x*-axis) and after (*y*-axis) clearing with SeeDB-Live. The comparison was performed as described previously ^63^. Orientation tuning was not affected by clearing with SeeDB-Live. n.s., non-significant (Wilcoxon signed-rank test). **(f)** Epifluorescence images of L2/3 neurons in S1 labeled with Voltron2_549_-ST in an anethetized mouse (P17) before and after clearing with SeeDB-Live (1 hour after clearing). **(g)** Action potentials were detected in a L2/3 neuron indicated as #1 in **(f)**. **(h)** Traces of the voltage changes in L2/3 neurons indicated in **(a)**. Ticks indicate detected action potentials. **(i)** Dendrites of a L2/3 neuron in S1 labeled with mTurquoise2 and Voltron2_549_. We imaged an awake mouse (2 months old) after SeeDB-Live treatment. Imaging depth: 90 µm. **(j)** Two-photon fluorescence image of mTurquoise2 (left) and epifluorescence image of Voltron2_549_ (right). A representative result is shown. **(k)** The spikes were detected at the dendrites of an L2/3 neuron in S1 of an awake mouse indicated in **(j).** The red ticks indicate the synchronous events that coincided among the dendrites.

**Supplementary Figure 10.**
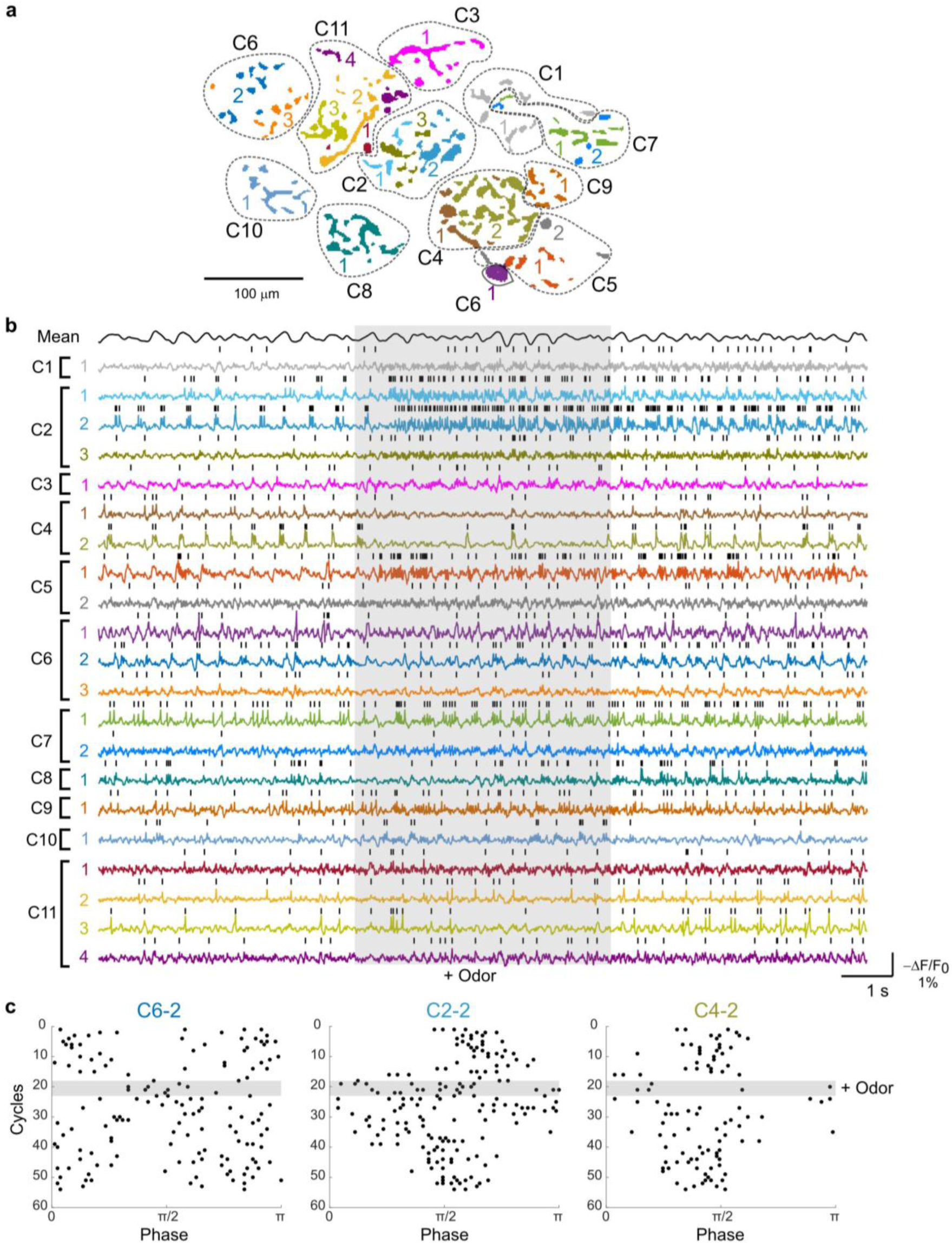
Widefield voltage imaging of mitral/tufted cell dendrites in the olfactory bulb *in vivo*. **(a)** Clusters (encircled by dotted lines) and subclusters (shown in different colors) of ROIs (mostly dendrites) of the mitral/tufted cells in the mouse olfactory bulb. The clusters and subclusters were defined based on voltage traces and spike synchronicity of ROIs, respectively. See also Supplementary Video 12 for Voltron2_549_ −ΔF/F_0_ images in ROIs. **(b)** Voltage traces of all subclusters (−ΔF/F_0_). The black trace on the top (Mean) shows the averaged ΔF/F_0_ of all glomeruli representing the theta wave in the olfactory bulb. Odor (1% amyl acetate) was delivered to the mice during the gray shaded period. Ticks indicate the detected action potentials. **(c)** The spike timing in each sniff/theta cycle. The timing is shown by the phase of the theta cycle. Odor (1% amyl acetate) was delivered to the mice in the gray shaded time. The peaks of the wave were regarded as π/2 phase.

